# Reliability of sucrose preference testing following short or no food and water deprivation - a Systematic Review and Meta–Analysis of rat models of chronic unpredictable stress

**DOI:** 10.1101/2023.02.22.529490

**Authors:** Jenny P. Berrio, Sara Hestehave, Otto Kalliokoski

## Abstract

The sucrose preference test is a popular test for anhedonia in the chronic unpredictable stress model of depression. Yet, the test does not always produce consistent results. Long food and water deprivation before the test, while often implemented, confounds the results by introducing unwanted drives in the form of hunger and thirst. We assessed the reliability of the test when only short or no fasting was used. We searched PubMed, Embase, and Web of Science for studies in rats exposed to chronic unpredictable stress that used no more than six hours of food and/or water deprivation before the test. Sweet consumptions, for stressed and control/antidepressant-treated animals, in 132 studies were pooled using random effects models. We found a decrease in sweet consumption in stressed rats, compared to controls, that was halved when a non-caloric sweetener was used and significantly reduced when sucrose consumption was corrected for body weight. What is more, the length of food and water deprivation was found to confound the effect. The effect was reversed when the stressed rats were treated with antidepressants. Methodological strategies meant to control for recognized sources of bias when conducting the test were often missing, and so was a clear and complete report of essential study information. Our results indicate that not only is food and water deprivation before the test unnecessary, but not recommended. Even in absence of long fasting, we found evidence of an additional effect on sweet consumption that is unrelated to anhedonia. Without properly controlling for non-hedonic drivers of consumption, the test is unreliable as a proxy measure of anhedonia. Strengthening the methodological rigor and addressing the confounding effect of metabolic factors in the sucrose preference test prevents misleading conclusions that harm the translatability of the associated research and perpetuates the use of animals for little gain.

## Introduction

The development of the chronic unpredictable stress (CUS) model of depression was based on the observation that exposing rats to a series of stressors (*e*.*g*., food and water deprivation, wet bedding, social crowding), over several weeks, caused a decrease in their consumption of a sweet solution^1,2^. This effect could be reversed by antidepressants^2^. The observed changes were assumed to be indicative of a reduced responsiveness to rewards, and hence a telltale sign of stress-induced anhedonia, a core symptom of depression^3^. Anhedonia is a state of decreased ability to feel pleasure and a reduced interest in activities that were previously found to be enjoyable. The preference for a sweet solution is assumed to be in proportion to the pleasure experienced in its consumption^4^, and so, when stressed rats consume less solution than do unstressed rats, it is suggested that they have comparatively less capacity to experience enjoyment. Thus, testing for sweet preference, using the so-called sucrose preference test (SPT), became widely accepted as a test for anhedonia.

Given its simplicity, the SPT became routine practice in validating the CUS model, and today it is hardly limited to this model. Other models of depression, induced using other forms of chronic stress, use it to assess reward sensitivity. It plays an important role in screening novel antidepressants and therapeutic strategies^5–7^. However, conflicting results have been reported throughout the years, casting doubts on the reliability of the CUS model and the SPT as an outcome measure^8–10^. To understand these concerns, a brief mention of the methodology of the test is warranted. Before being tested, the animals are commonly fasted. The length of this food and/or water deprivation varies, but usually more than 12 hours are employed^11,12^. The rationale for the fasting is tied to length of the test. Per tradition, once the animals are presented with either one bottle containing the sweet solution, or more often, two bottles (water and sweet solution), only one hour is allowed for consumption^2^. Thus, long fasting is used to motivate the animals to consume considerable amounts within a short time. Once the test is over, either the consumed amount of sweet solution, or its preference over water, a calculated measure that accounts for variations in the total amount of consumed liquid, is registered. Stressed animals are then compared to controls to infer their hedonic state.

Ten years after the conception of the CUS model, two papers^13–15^ highlighted the influence of food and water deprivation and the nutritional state of the tested animals on the outcome of the SPT. A correlation was observed between the decrease in sucrose consumption in stressed animals and retarded body weight gain as a result of the exposure to stress^14^. The effect of stress on the SPT results was abolished after a simple correction for body weight (as expressed by the authors: “the bigger the rat, the bigger the consumer”). What is more, similar artifacts in the SPT results were noted after periodic food and water deprivations alone (without any additional stressors)^13^. Similarly, studies substituting sucrose in the sweet solutions for the non-caloric saccharin, found that results were dependent on hydration^16^ or the nutritional status of the animals^17^. Food deprivation was necessary to elicit reduced saccharin intake in chronically stressed rats^17^. Whereas 24 hours of water deprivation resulted in a decreased preference, 5-8 hours did not^16^. In a series of papers aimed at easing these concerns and reaffirming the premise that CUS causes a decreased sensitivity to rewards^10,18^, Paul Willner, the originator of the model, made some concessions. The stressful component of fasting procedures, and their effects on body weight, may influence the test. Willner recognizes weight loss as a possible confounder and grants that sweet consumption could be influenced by external factors such as nutritional state. Despite the general agreement on the confounding effect metabolic factors could have on the SPT, long periods of food and water deprivation continue to be used as a stressor and as an incentivizer. A recent user survey^11^ found that 83-86% of CUS protocols in rats use food and water deprivations as a stressor, and 32 out of 57 labs use more than 12 hours of fasting to incentivize consumption in the SPT.

Food or water deprivation, while stressful^19^, are not necessary to make rodents consume a sweetened solution. Rodents readily consume non-caloric sweeteners over water^20^. Thus, any anhedonic effect brought on by CUS should be measurable also when no food and water deprivation is performed before a SPT. In a test that measures the consumption of a caloric solution by a stressed (often underweight) rodent, thirst and hunger are bound to confound the interpretation of the results. When offering a choice between water and a sweet solution, one may wonder if thirst would result in increased consumption of whichever liquid the animal approached first, rather than it being a case of preference. Hunger leaves no doubt, it clearly drives the consumption of caloric sweet solutions. Because other uncontrolled factors beyond the pleasure experience are influencing the consumption, variable responses can be observed in the SPT and less accurate conclusions can be drawn. Thus sweet consumption is rendered less valid as a putative measure of anhedonia as the readout is not free from metabolic artifacts^15^. Eliminating the practice of fasting the animals before the test is not only likely to improve reproducibility but will also align with the 3R principles of humane research^21^.

Our study aimed to assess the reliability of the SPT when no fasting, or only short periods of food and/or water deprivation, were used before the test. We conducted a systematic review of studies in rats exposed to CUS that were assessed in the SPT after being fasted for 6 hours or less. We hypothesized that stressed animals should show a decreased sweet consumption, compared to controls. This decrease should persists when a non-caloric sweetener is employed (saccharin), and when the outcome measure is expressed as total liquid intake and corrected for body weight. Likewise, chronic treatment with approved antidepressants should show a reversal/improvement of the effect in treated rats (increased sweet consumption) compared to stressed conspecifics.

## Methods

### Registration and open access data

The protocol was registered in PROSPERO (CRD42021259015). We followed the PRISMA guidelines for reporting^22^. Detailed information on the methods, as well as links to the data repository, can be found in the supplementary materials. Deviations from the protocol are listed in the supplemental materials.

### Search strategy

For clarity purposes, the term “sucrose preference test” (SPT) is used throughout the article to refer to tests employing both sucrose and saccharin. Publications from 1987 to June 2021 were retrieved from three databases: Medline via PubMed, Embase and Web of Science. In all databases, three search strings were created and combined. Each string searched in the title and abstract for studies in (1) rats, (2) using CUS and (3) the SPT.

### Eligibility criteria

After removal of duplicates, two reviewers screened the studies independently in two phases. An initial title and abstract screening performed using the web-app Rayyan^23^ excluded studies that: (a) did not describe an original study in laboratory rats, (b) used a stress model other than the CUS and, (c) did not assess anhedonia. Whenever the reviewers disagreed, or there was incomplete information for an exclusion, the study was included in the next phase. A second screening was performed on full texts. During this phase, only studies that used CUS for at least two weeks in weaned rats, and that used the SPT to assess anhedonia, were included. The studies needed to report on a familiarization phase to the sweet solution and no more than 6 hours of food/water deprivation could be employed. Moreover, the SPT results in a cohort of stressed animals had to be compared to those of a cohort of unstressed animals (controls), or a cohort treated with an FDA-approved antidepressant for at least 2 weeks while still undergoing stress. A third reviewer resolved discrepancies at this second stage. For a complete description of the criteria used at each stage, see Supplementary table 1.

### Study details and outcome data extraction

One reviewer extracted the methodological details from each study (Supplementary table 2A). The outcome extraction of all included experiments was performed by two reviewers, independently. Sweet consumption at the end of CUS, for stressed and control/drug-treated groups, was extracted (Supplementary table 2B). During this stage, if there were more than one experiment in a study that fulfilled the inclusion criteria, each experiment was extracted individually. We report the number of experiments unless otherwise specified. When a study reported the outcome of the same cohort of animals in more than one measuring unit (*e*.*g*., preference and intake) or in response to different sweets, these alternative outcomes were also extracted. The calculated sweet preference was our main outcome of interest (Supplementary table 3). The alternative outcomes were used in subgroup analyses only. Data were preferentially extracted from raw data, tables or text. When results were exclusively presented in figures, data were extracted using a digitizing tool^24^. Discrepancies in the outcome extractions were resolved by the reviewers reaching a consensus decision. Efforts were made to contact authors in the case of incomplete data. If the data could still not be obtained, the experiment was excluded. To explore the effect of the length of food and water deprivation on sweet consumption, 10% of the studies that were excluded during the full text screening because fasting exceeded 6 hours were randomly selected. Their study details and sweet consumption at the end of CUS for the stressed and control groups were extracted in a similar fashion to the included studies (see further details in supplementary material).

### Quality and risk of bias assessment

The quality of reporting and the internal validity was assessed independently by two reviewers, while a third reviewer resolved discrepancies. A checklist adapted from the ARRIVE essential 10^25^ was used to assess the quality of reporting in 100 studies randomly selected from the pool that fulfilled all inclusion criteria (Supplementary table 4). A checklist adapted from SYRCLE’s risk of bias tool^26^ was used to assess the methodological quality of all 132 studies included in the meta-analysis. Eleven items assessing selection, performance, detection, attrition and reporting biases were checked for each study (Supplementary table 5). Each study was assigned a risk of bias score totaling the items in the checklist that had clearly been addressed.

### Data synthesis

Statistical analyses and visualizations were done in RStudio 2022.02.1.461^27^ with R version 4.2.1^28^. For a complete list of the packages used, see the supplementary material.

A meta-analysis was deemed feasible if at least 10 studies were included. Pooling and weighting of individual effect sizes (standardized mean differences, SMD - Hedges’ g) in the SPT was done with a random-effects model using the DerSimonian and Laird method (DL) for estimating between-experiment heterogeneity. Separate pooled effects were calculated for stress and antidepressant treatment, respectively. Heterogeneity was estimated using the I^2^ statistic^29^. Prediction intervals were used as a measure of precision. Studies were not excluded on the basis of the risk of bias. Subgroup analyses were used to test for differences in the pooled effect when the caloric content of the test solution (sucrose *versus* saccharin), total liquid intake (intake *versus* preference) and body weight (intake corrected for body weight *versus* preference) were taken into account. Pairwise analyses were conducted for experiments that reported preference and intake, or preference and intake corrected by body weight, in the same cohorts of animals. For these analyses, the pooled effect of the difference in effect sizes between stressed animals and controls for both outcome measures was estimated using a fixed effects model. The sample size for both outcomes was calculated as the sum of the sample size in the stressed and control groups divided by two. The effects of the strain, sex, age, photoperiod of the test, and the class of antidepressant were explored in subgroup analyses, while the effects of the length of food and/or water deprivation before the test, length of stress, length of antidepressant treatment, and risk of bias score were evaluated using univariate meta-regressions. The subgroup analyses were performed when there were more than four studies per variable of interest. A meta-regression was employed to examine if the effect size of a study depended on the length of deprivation. Studies using more than 6 hours of food or water deprivation were included in this analysis. Sensitivity analyses exploring robustness of our models to the effects of sample size inaccuracy (due to poor reporting) and choice of heterogeneity estimator were performed (supplementary material).

### Publication bias

For studies that reported results as a preference we constructed a funnel plot to assess whether there was an artificial inflation of the estimated effect of stress on the SPT. For an unbiased assessment, visual scrutiny was complemented by Egger’s test. Small study effects, including publication bias, were assessed for all experiments, including those assessing the response to antidepressants, and for subgroups with more than 10 studies (Supplementary material).

## Results

### Selection process

3,277 reports were found through the combined search in three databases. After removing duplicates and screening titles and abstracts, 1,035 reports were found to describe an original study on laboratory rats that underwent CUS and were assessed for anhedonia. After full-text screening, 876 reports were excluded because they did not fulfill our criteria (Figure 1); 81.8% of the studies in rats exposed to CUS were excluded because they employed food and/or water deprivation longer than 6 hours before the test (Figure 2). 159 studies fulfilled all the inclusion criteria. 27 were excluded after outcome extraction, most because of incomplete data. A total of 132 publications, with a total of 183 unique experiments were included for analysis.

**Figure 1.**
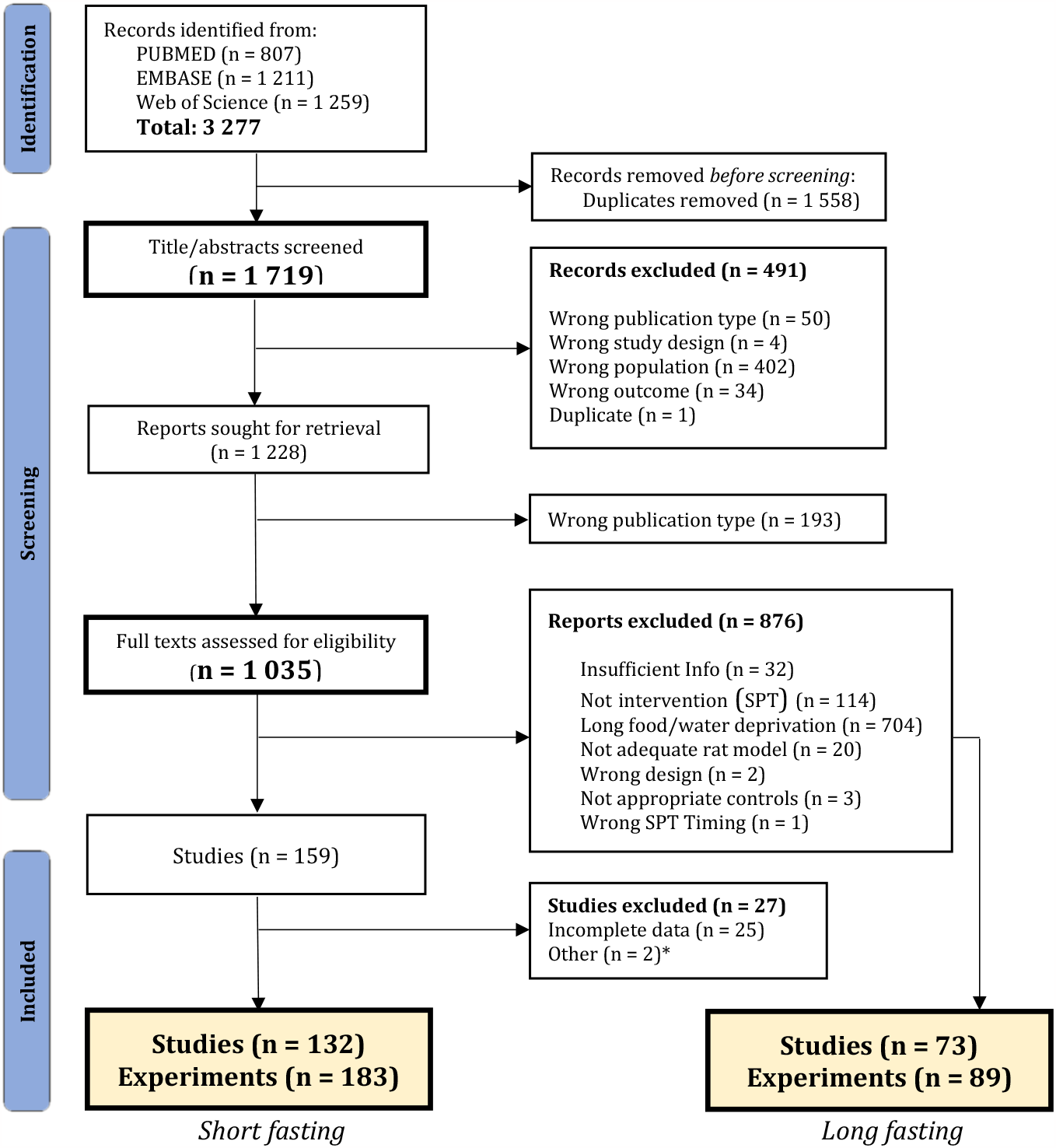
Systematic search flow diagram. Summary of the selection process during both the title and abstract screening and the full-text screenings. For a detailed description of the exclusion criteria see Supplementary table 1. From an initial sample of 1,719 unique records, 159 fulfilled all inclusion criteria after full-text screening. Of those, 132 fulfilled all criteria and had complete data. 183 unique experiments were extracted from those studies and were included in the meta-analysis. *1. Russian translation of another included publication; 2. Study was excluded because of an intervention administered concurrently to the CUS. Seventy three studies that employed long (> 6 h) periods of food/water deprivation were randomly selected to evaluate the influence of length of fasting in sweet consumption. Adapted from the “PRISMA 2020 flow diagram for new systematic reviews”^22^.

**Figure 2.**
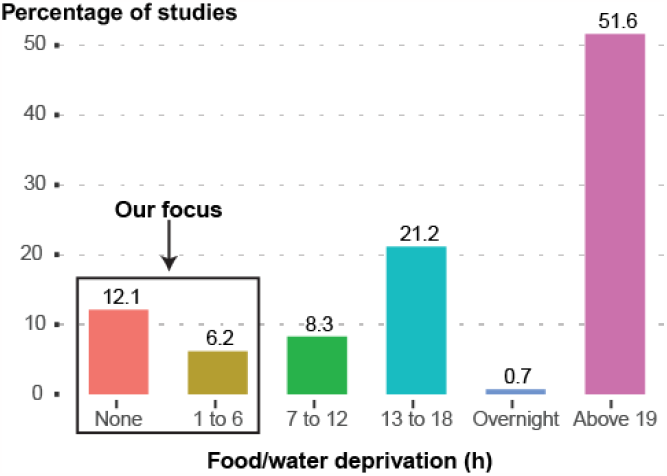
Percentage of screened studies that performed SPT by length of food and water deprivation before the test. The studies graphed include those employing rats exposed to CUS that habituated to the sweet and performed the SPT. This includes studies that were either included in our analysis or excluded because of other criteria (860 studies: 728 excluded, 132 included). 73.5% employed more than 12 hours of fasting, with the biggest proportion (51.6%) employing more than 19 hours of fasting. Only 12.1% of the screened studies did not deny animals food or water before the test.

### Characteristics of the included experiments

#### Model characteristics

Typically, researchers used male Sprague Dawley rats, 9 weeks of age or less, that were stressed for 2 to 4 weeks (Figure 3). Only 5.5% (10 experiments) used females (Figure 3C). Wistar rats were the second most common choice (30.1%/55 experiments) (Figure 3A). Only a small proportion of experiments performed more than 6 weeks of stress (11.4%/21 experiments) (Figure 3D).

**Figure 3.**
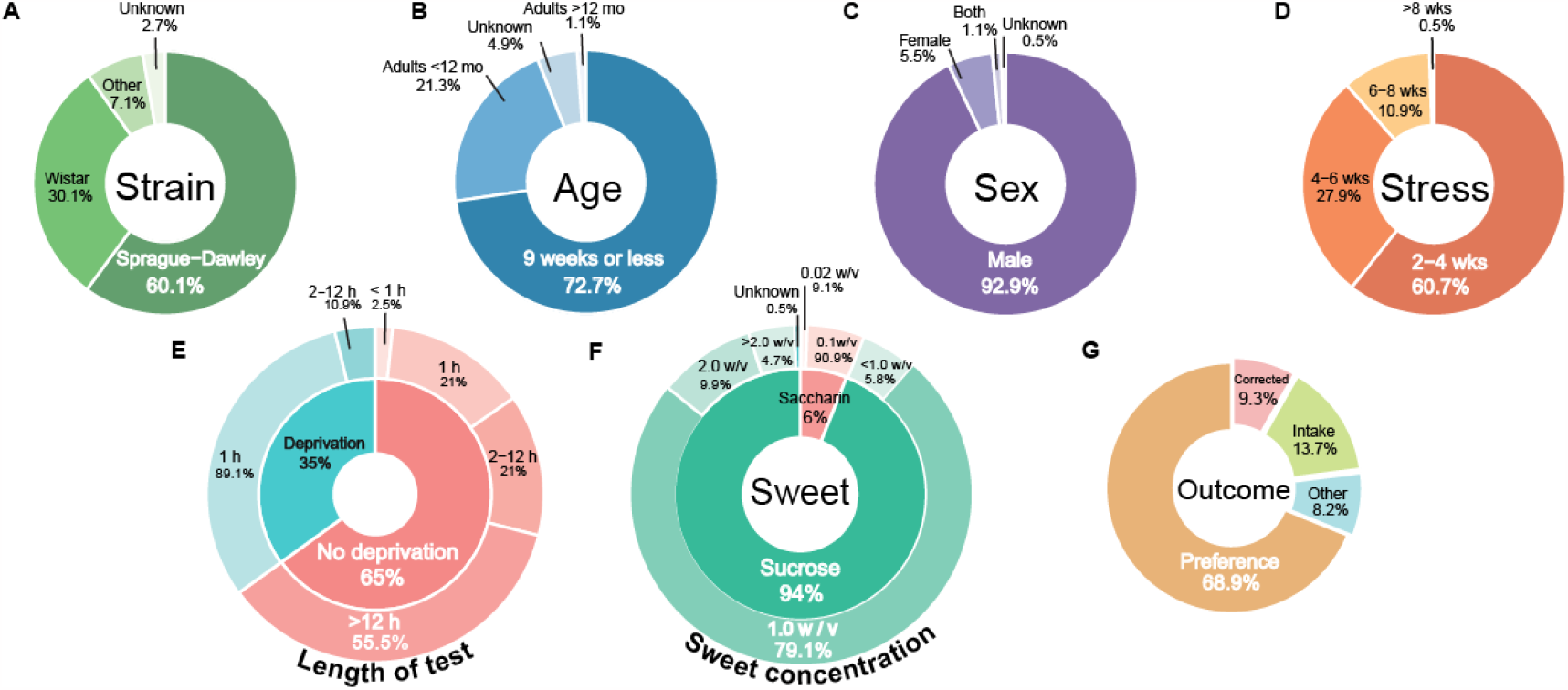
Model characteristics of the included experiments. Distribution of the included experiments by (A) rat stock/strain, (B) sex of the animals, (C) age and (D) length of the chronic unpredictable stress in weeks. *Methodology of the SPT in the included experiments*. Charts of (E) food and water deprivation (inner ring) and the length of the SPT that followed it (outer ring), (F) the type of sweet (inner ring) and the sweet concentrations employed (outer ring), and (G) the reported outcome. “Preference” is calculated as the consumption of sucrose solution relative to the total liquid consumed. “Corrected” is used to represent the measures that divided the individual sweet intake (g or mL) by body weight. Total of 183 experiments. Abbreviations: mo, months, wks, weeks.

#### SPT Methodology

The majority of experiments used two bottles during the SPT, 17 experiments (9.3%) presented the rats with only one bottle containing the sweet solution. Sucrose was the most common type of sweetener (93.9%/172 experiments) (Figure 3F - inner ring), with concentrations that ranged from 0.2% to 30%. A 1% sucrose solution was by far the most common concentration (Figure 3F - outer ring). All but one experiment employing saccharin used a concentration of 0.1% (Figure 3F). In 65% (119 experiments) of the experiments rats were not deprived of either food or water before the SPT (Figure 3E - inner ring). The length of the test ranged from 15 minutes to 8 days. Around half of the experiments using no food or water deprivation performed tests longer than 12 h (55.5%/66 experiments)(Figure 3E - outer ring), while 21% (25 experiments) used the typical 1-hour test. By contrast, 57 of the 64 experiments (89.1%) denying the animals food and/or water used a 1-hour test (Figure 3E). The test results were most commonly reported as a percentage of intake (preference) (68.9%/126 experiments), followed by absolute intake in weight or volume (13.7%/25 experiments). Only 9.3% (17 experiments) corrected the outcome for body weight (Figure 3G). 34% of the experiments with tests shorter than 12 hours carried out the test during the animals’ active phase (the dark photoperiod) of the day, while in 50% of these experiments the photoperiod was unknown (Supplementary figure 1).

#### Antidepressant treatment

All twelve experiments exploring antidepressant treatments in stressed rats used sucrose (Supplementary material). Experiments were performed in male rats, either Wistars (7 experiments) or Sprague Dawleys (4 experiments). Only one study did not report the strain. The majority of protocols (9 experiments) did not employ food or water deprivation before the test, only 3 experiments employed 4 hours of fasting. The length of the stress protocol before the antidepressant treatments ranged from 2 to 6 weeks with an average of 3.3 weeks. Likewise, the length of the antidepressant treatment ranged from 2 to 5 weeks, with an average of 3.1 weeks. The two most common antidepressants were fluoxetine (typically 5 or 10 mg/kg) and imipramine (10 or 20 mg/kg). The other antidepressants were venlafaxine, clomipramine, and escitalopram.

### Quality of reporting

Eighteen of the 100 studies randomly selected for assessments of reporting quality were not included in the meta-analysis, therefore only 82 studies were analyzed (Supplementary figure 2). Most of the studies presented a description of the animals and of the study design, and reported the outcome measures and the statistical methods used to analyze them (79-95%). Less than half of the studies clearly reported the strategy to allocate the experimental units, presented a full description of the stress protocol or gave a complete report of the results of the SPT (46-48%). Less than a third of the studies were clear about the exact numbers of rats included in the statistical analysis and explained any exclusions (27%). A clear statement of the sample size with its calculation and any reference to blinding was present in only 5% of the studies. No study reported all items.

### Risk of bias

The 132 included studies were assessed for risks of bias. On average, 2.9 items out of 11 had low risk of bias. Most of the studies had a substantial number of unclear risks because of poor reporting of essential information (Supplementary figure 3). Notably, only about 19% of the studies reported that the experimental groups were balanced in key characteristics at baseline (sex -in studies where both were used - weight, and baseline sweet preference). For most experiments, this was unclear (64%). In 21% of the studies, the control and stress groups were housed separately under similar environmental conditions to avoid the indirect exposure of controls to stress. 16% of studies counterbalanced the bottles (*i*.*e*., swapped the physical locations of the sweet solution and water bottles in the cage) within the test period to avoid place preference. 17% of the studies were classified as high risk in this item because they either used only one bottle during the test or performed no counterbalancing within of the testing period. Around half of the studies (55%) had a low risk of reporting bias and a third (33%) had a low risk of attrition bias. 9.1% of the studies had unexplained missing data.

### Data synthesis

#### Sucrose preference test in stressed rats

Stressed rats were compared to controls in 171 experiments. On average, chronically stressed rats had a significantly reduced consumption of sweet solution compared to unstressed controls (SMD -1.44; 95% CI: -1.61, -1.27; p < 0.0001)(Figure 4A). However, high variation in the results was observed (between-experiment heterogeneity: I^2^ = 79.3%; 95% CI: 76%, 82%).

**Figure 4.**
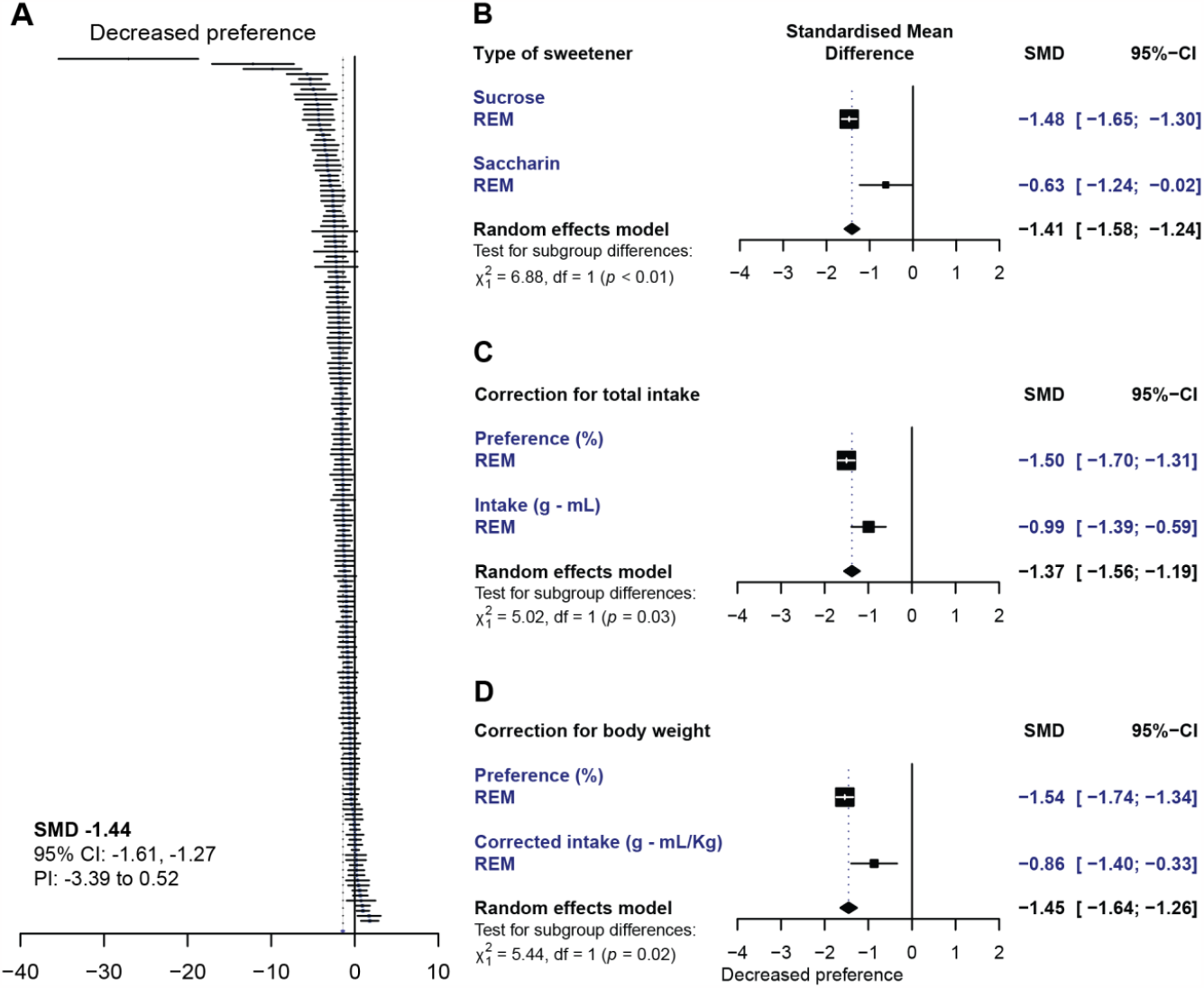
*A*. Simplified forest plot of the sweet consumption in stressed animals compared to unstressed controls. Each line represents the outcome of an individual experiment with its 95% confidence interval. The dotted line to the left of the zero line represents the average effect estimate.

##### Caloric content of the solution

Experiments that used sucrose (160) were compared to those employing saccharin (14) to test for differences in the results dependent on the caloric content of the solution. Three experiments measured the response to both sucrose and saccharin in the same cohort of animals (Figure 4B). Although stressed rats consumed less sweet solution compared to controls regardless of the type of sweetener, the magnitude of the effect was significantly lower in experiments using saccharin (p = 0.0085). High between-experiment heterogeneity was present in both subgroups of experiments (sucrose: I^2^ = 78%; saccharin: I^2^ = 84%).

##### Total liquid Intake

Experiments reporting preference were compared to those reporting sweet intake. A total of 158 experiments that presented two bottles to the animals during the test were included in the analysis: 117 reported preference as the outcome, while 41 reported intake. Less consumption of the sweet solution was found in stressed rats compared to unstressed rats when either outcome was reported (Figure 4C). However, the magnitude of the effect was larger when reported as a preference (p = 0.03). High between-experiment heterogeneity was present in both subgroups (preference: I^2^ = 75%; intake: I^2^ = 82%). In 22 experiments both measures were reported for the same cohorts of animals. When these experiments were analyzed, a difference between the outcomes was not observed (pairwise comparison, fixed effects model: SMD 0.03; 95% CI: -0.21, 0.26; p = 0.83; Supplementary figure 5).

**Figure 5.**
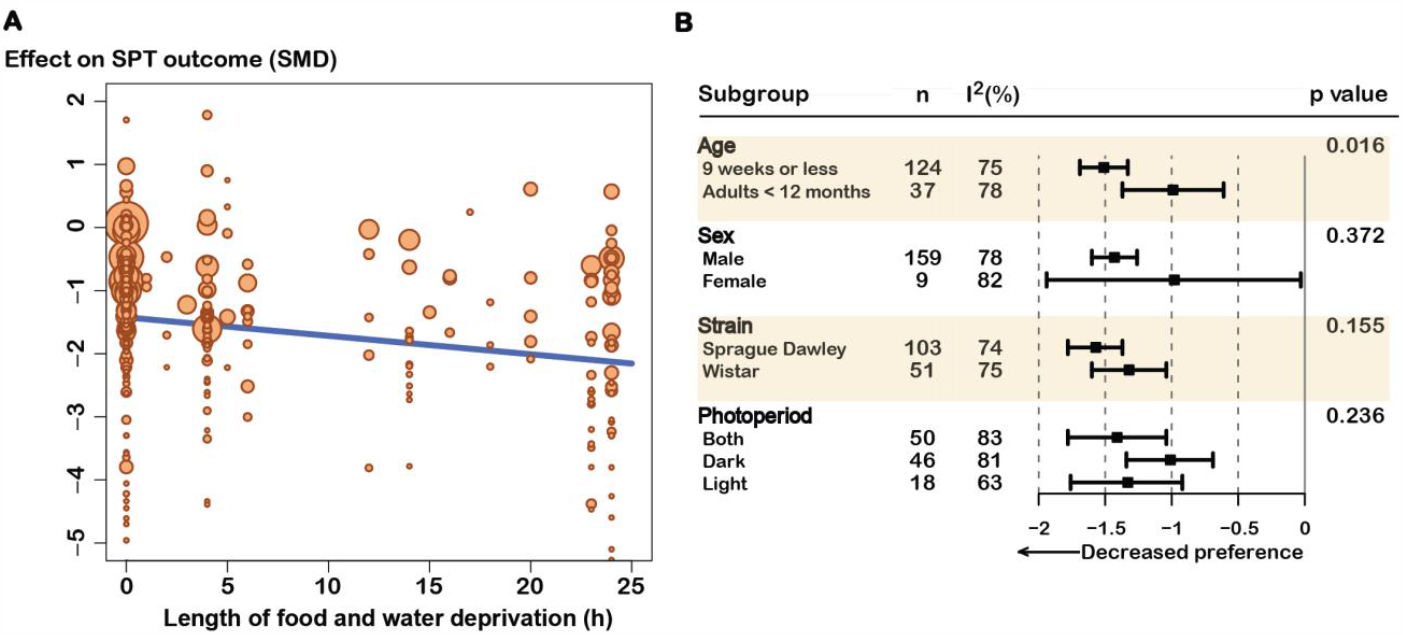
A. Bubble plot displaying the result of the meta-regression of length of food and water deprivation when studies using short fasting and long fasting were included. Each symbol represents an individual experiment and its size the inverse variance of the estimated effect (the bigger the symbol, the smaller the variance). Some extreme effect values excluded from the figure to favor visualization of the relationship. The results are presented in Supplementary table 6. *B. Subgroup analysis of categorical variables of interest*. Only variables that had more than 4 independent studies were included in the analysis. The n represents the number of experiments included in each subgroup. The p value concerns the test for subgroup differences. Abbreviations: SMD, standardized mean difference; I^2^, between-experiment heterogeneity.

##### Body weight

Experiments reporting on sucrose preference (112 experiments) were compared to those reporting the intake of sucrose corrected for body weight (g/kg or mL/kg)(18 experiments). Both measures showed a decreased sucrose consumption in stressed animals. However, the magnitude of the effect was significantly lower (p = 0.02) when the intake was corrected for body weight (Figure 4D). High between-experiment heterogeneity was present in both subgroups (preference: I^2^ = 75%; corrected intake: I^2^ = 81%). Five experiments (4 studies) reported the results in both ways for the same animals. When these experiments were analyzed, a difference between the outcomes was not observed (pairwise comparison, fixed effects model: SMD 0.00; 95% CI: -0.32, 0.32; p = 1.0; Supplementary figure 6).

Studies falling on the left side of the zero line found a decreased consumption in stressed animals. For a more detailed forest plot see Supplementary figure 4. Forest plots of subgroup analyses comparing (B) studies using sucrose *versus* those using saccharin (Type of sweetener), (*C*) comparing those that accounted for total liquid intake by calculating the percentage (%) of preference (sweet solution intake relative to total liquid intake) and those reporting intake (in grams or milliliters), and finally, (*D*) comparing those that reported sucrose preference versus those reporting intake corrected for body weight. Abbreviations: SMD, standardized mean difference; 95%-CI, 95% confidence interval; PI, prediction interval.

##### Length of fasting

The length of food and/or water deprivation before a SPT was a significant predictor of the sweet consumption. Longer periods of fasting were associated with larger effects. We observed that the reported sweet consumption among stressed rats compared to controls significantly decreased by 0.03 standard deviations per hour of deprivation; that is, 0.72 standard deviations after a day of fasting (p = 0.0003)(Figure 5B and Supplementary table 6). Notably, this outcome remained consistent even after considering the type of sweet and accounting for the duration of stress (supplementary material). However, this finding explained only 3.16% of the between-experiment variation in the results.

##### Other sources of variability

Likewise, sex, strain/stock or the period of day when the test was performed did not significantly change the magnitude of the effect. Animals of 9 weeks of age or less had a significantly greater reduction in sweet consumption from stress than did older rats (Figure 5B). With increasing length of stress, the sweet consumption decreased in stressed rats relative to controls by 0.13 standard deviations per week. However, only a fraction (1.68%) of the between-experiment variance was explained by this variable (Supplementary table 6). The risk of bias score was also not a significant predictor. Heterogeneity was largely unaffected by these predictors.

#### Sucrose preference test in rats treated with antidepressants

The response to antidepressant treatment for 2 weeks or more was assessed in twelve experiments. Overall, stressed rats treated with antidepressants had a significantly increased consumption compared to stressed controls (SMD 1.85; 95% CI: 1.14, 2.56; p < 0.0001)(Supplementary figure 7). High between-experiment heterogeneity was observed (I^2^ = 74.6%). Neither the duration of fasting, the length of the stress protocol or antidepressant treatment, nor the risk of bias score were significant predictors of the SPT results (Supplementary table 6). Likewise, neither strain/stock (Sprague Dawley or Wistar) nor class of antidepressant (SSRI or tricyclic antidepressant) significantly changed the magnitude of the effect (Supplementary figure 8 and 9, respectively).

### Funnel plot analysis

Since formal analysis of funnel plots using SMD are known to be susceptible to bias, we applied Egger’s regression test to the subset of tests reporting their results as a preference. Visual assessment of the funnel plot (Figures 6A) revealed some asymmetry with respect to the studies comparing stressed rats to controls. Egger’s regression confirmed the asymmetry (intercept: -3.77, t: -5.98, p < 0.0001). Similar results were seen for subgroup analyses (Supplementary material).

**Figure 6.**
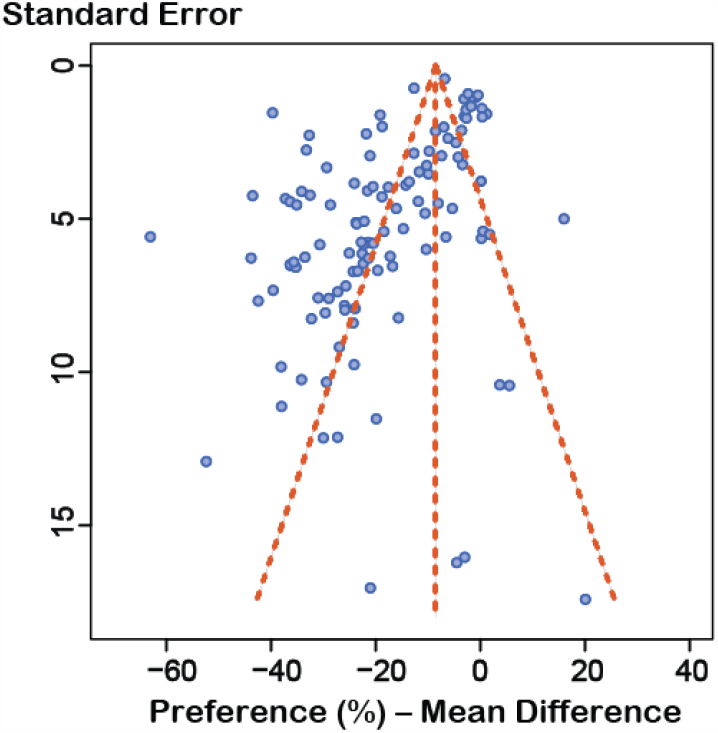
Funnel plot of experiments reporting preference (n = 117). The standard error of the experiments is plotted against their observed effect sizes (MD). The y-axis is inverted with studies with the lowest standard errors on top.

## Discussion

Despite the widespread use of the SPT, failed attempts to reproduce results are still prevalent. Many biological and methodological factors have been hypothesized to underlie these inconsistent results^9^. Among them, performing long food and water deprivation before the test has been recognized as an important confounder that contributes undesired metabolic noise. Rodents do not need to be hungry and thirsty to consume sweet foodstuff^20,30^. Under the premise that sweet taste is rewarding, that CUS changes the experience of pleasure derived from its consumption, and that the SPT is a valid measure of such an experience, the SPT should detect a change in the rewarding properties of the sweet solution (a decreased consumption) in stressed rats also when food and water are available. Indeed, this is the only scenario where one can be confident the observed changes relate to the hedonic value of the sweet, rather than to underlying metabolic drives. Yet, long food and water deprivations are performed routinely. Over 70% of the studies that were screened in our study denied rats of food and/or water for more than 12 hours before the test, a percentage that is comparable to previous findings^11,12^.

Why is such practice still common among researchers despite the growing recognition of its confounding effect? Tradition might be at fault. Long food and water deprivations are traditionally used to motivate rodents to consume considerable amounts of sweet solution in short periods of time, usually within an hour^2^. But if tradition comes at the cost of validity, perhaps it is time to challenge tradition. A recent meta-analysis investigated the reliability of the most commonly used behavioral paradigms used to test rodents modelling anxiety.

Most tests unfortunately had low to no sensitivity to anxiolytics used to treat anxiety in humans, and poor predictive validity for the discovery of new treatments. What is more, the results of the tests were so heterogeneous and often contradictory, that the authors advocated for a re-evaluation of the validity of the tests^31^. With our study we aimed to assess how reliably the SPT finds a decreased consumption when no more than 6 hours of food and water deprivation are used before the test. We found an overall decrease in sweet consumption in stressed rats, compared to controls. Although there was a significant variability in results across different experiments. A beneficial effect of antidepressant treatment was also observed. However, we only identified twelve studies that measured this response. Importantly, we found that not only is food and/or water deprivation unnecessary, it is inadvisable. We were able to find more than a hundred published experiments where the SPT was carried out without any previous food or water deprivation, and with overall positive results. What is more, we found that longer periods of fasting were associated with larger effects. However, this does not mean that longer deprivations strengthen the outcome of the model. Instead, each hour of deprivation artificially inflates the observed effect, independently of stress, distorting the results and creating an overly optimistic picture of model’s success. Indeed, a week’s worth of stress (0.13 SD) is overshadowed by the confounding effect of as little as 5 hours of fasting (0.15 SD).

The decrease in sweet consumption in response to chronic stress was still present when a non-caloric sweet solution was employed, and when outcome measures that correct for total liquid intake and weight were used. However, while stressed animals still consumed less saccharin than did controls, we found that the magnitude of this difference was more than halved compared to experiments using sucrose. This change in effect is to not be ignored. This may be explained by discrepancies of the caloric component of the sweetener; or alternatively, methodological factors like concentration-selections, where the traditional saccharin-concentration could lead to more robust preference, and thereby less sensitive to modulation by stress. Granting the cause is open to inquiry, this finding comes with an additional warning of undesired non-hedonic noise, even when long deprivations before the test are not employed. Stress protocols often use periodic fasting as a chronic stressor^11^. These food and/or water deprivations not only have shown to cause decreased sweet consumption on its own^13^, but they are also associated with changes in weight, metabolism and the mechanisms controlling appetite of rats^32,33^. In our study, when sweet consumption was corrected for body weight, the effect of stress on the SPT results was also attenuated. Although the paired analysis of four studies could not substantiate this finding, observing significant changes in the results of the test when a non-caloric sweetener is used and when the consumption of a caloric solution is corrected for body weight should caution researchers against regarding the results of the test as a reliable measure of anhedonia. Unfortunately, only a small number of the analyzed experiments employed saccharin, or accounted for body weight, showing that most researchers employ sucrose and seldom correct for the possible confounding effect of weight in their model. Methodological safeguards designed to minimize bias while performing the test were also often missing. Since considerable inter-individual variability in drinking patterns and sweet preference can be observed^7,30,34^, and differences in body weight inherently introduce metabolic heterogeneity, ensuring the animals are balanced across experimental cohorts in these aspects at baseline is important^34^. Yet, in only 19% of studies such balancing was reported.

Likewise, counterbalancing the bottles within the test period to avoid a strong preference for bottles placed on a particular side of the test cage, a recognized confounder of the SPT in rats and mice^7,34^, was reported in only 16% of the studies. What is more, avoiding indirect stress affecting the control rats through olfactory, visual, and auditory signals, an effect that has been demonstrated in rodents^35–37^, was clearly ensured in only 21% of the studies.

This lack of methodological rigor in properly controlling for non-hedonic drivers of consumption clearly decreases the confidence in the results of the test as a measure of anhedonia (*i*.*e*., the internal validity). Under these circumstances, one cannot confidently relate a change in sweet consumption in the SPT to a change in the rat’s reward system. In the scenario where such a change is indeed caused by a change in the likeability of the sweet, isolating it and quantifying its magnitude becomes impossible when other factors that modulate consumption are eclipsing it. Thus misleading conclusions are reached that risk painting an inflated and unrealistic picture of the change in reward sensitivity. Especially when sucrose preference test results are used to discriminate between resilient and susceptible animals, a practice that in turn overestimates the effect, thus escalating the problem as demonstrated in our follow up study^38^. This has tremendous implications for the translatability and ethics of the research in the area^39^. Not only are the effects that future studies base their power analysis on inexact, perpetuating the unethical cycle of animal use for inconclusive research, they also lead to overly optimistic predictions of the effects of new therapeutic strategies. Therapies that may fail when transitioning to human trials, wasting resources and time in the process. In order to increase the robustness of the SPT as a measure of CUS-induced anhedonia, it is paramount to completely abolish any food and water deprivation before conducting it. We also need to systematically incorporate methodological safeguards that minimize bias when measuring consumption in future studies.

In alignment with previous findings^12^, highly heterogeneous results between studies were observed. There was also considerable heterogeneity with respect to the ways in which the test was conducted. Differences in the SPT protocol are likely to be an important source of variation in the results across different studies and research groups. The most frequent design in our dataset – a two-bottle choice paradigm, performing no fasting, and measuring the consumption in a period of at least 12 hours, preferably during the dark photoperiod - would serve as a good frame for a standardized protocol moving forward. By deviating from the traditional 1-hour test^40^, increasing the length of the test, the differences from measuring consumption in different times of the light/dark cycle can be reduced. Rats are nocturnal and do most of their drinking during the dark phase^41^. While we saw no effect by photoperiod of the test, covering such critical period with longer tests should be a better measure of consummatory behavior in response to sweets^7^.

We found that strain/stock and sex, previously suggested to contribute significantly to the heterogeneity of SPT results^9,11,42^, did not have a clear effect on the studied experiments. Only age was associated with a difference in the SPT results. Sprague Dawley and Wistar rats, the most frequently used stocks, had similar decreases in sweet consumption and responded equally well to antidepressant treatment. This is similar to the results of a previous study^12^ that, although it found larger effect sizes in both stocks, did not observe significant differences. We found, as others have before us^5,12^, that most experiments used male rats exclusively. By contrast, women are more likely to suffer from depression than are men; they report greater severity of the depression, and are more likely to experience atypical symptoms or comorbid anxiety^43^. Yet, in our analyses, we cannot confidently make any conclusions with regards to sex-related differences in the response to the CUS model because the results from the few experiments employing females were highly heterogeneous. A similar case can be made for age. In humans, depression can occur at any point in life but the prevalence varies across life stages^44,45^. In our study, most experiments used rats that were at most 9 weeks old in the beginning of their study, a period of developmental transition for rats, from puberty to adulthood, thus making the subjects both “adolescents” and “young adults”^46^. When we compared these studies to experiments in older rats, we found a significant difference in the measured effect. Younger rats appeared to respond with a greater decrease in sweet preference following the CUS protocol. However, our numbers are somewhat inexact since we had to use body weight as surrogate marker for age in a number of experiments with poor reporting. Future studies would do well to clearly separate key developmental stages such as adolescence and adulthood when researching the role of age in stress related depression. Another important element of the model that we did not fully explore in our study is the stress protocol. We found that increasing the length of the stress was associated with larger decreases in consumption in stressed rats compared to controls, but we did not further analyze the details of such protocols. The types of stressors and their schedules vary greatly among research groups^11^ and so can their severity. Some choose to implement periods of fasting while others do not. The extent to which such divergent protocols are comparable and their relevance for modelling depression merits further validation^39^.

Unfortunately, most studies were assigned with an unclear risk of bias for most items due to the lack of clarity in their reports where important methodological information was often missing. In fact, we were unable to find a single study that reported on all of the items summarized in the ARRIVE essential 10 checklist^25^. We are not the first to report on this. Substantial risks of bias due to poor reporting have also been found by others^31^. This clearly has an impact on the robustness of meta-analytical studies such as the present one. Only a clear and detailed report allows other researchers to clearly understand the scope of a study, and more importantly, its limitations. What is more, publishing findings also when the results are unexpected increases the overall confidence in the evidence provided. We found evidence of a significant small study effect^47^. Whereas we cannot tell whether this stems from publication bias or simply biased study designs, we have a suspicion. Over the years we have met researchers telling us about their difficulties in obtaining expected results using the SPT, suggesting it is probable “unsuccessful” studies may have remained unpublished. Transparent reporting of studies, irrespective of outcome, is highly encouraged.

## Conclusions

In summary, we would like to urge caution with respect to our findings. While the SPT found a decreased sweet consumption when no fasting precedes it, we observed a significant change in the magnitude of the effect with longer periods of fasting, when a non-caloric sweetener was used and when sucrose consumption was corrected for body weight. This suggests that the observed response could have been influenced, at least partially, by underlying metabolic drives. We also found evidence of a lack of rigor in controlling for recognized confounders when conducting the test and a pronounced lack of clear reporting of essential methodological information. Without properly controlling for factors not related to the pleasure experience of consuming sweets, the outcome of the test in an unreliable proxy of changes in the reward system. Particularly if we consider that the model itself affects body weight and metabolism of the stressed rats. Thus, not only should the practice of fasting the animals before the test be eliminated altogether, but additional tuning of the test and the model might be required in individual studies. Albeit we only analyzed data from rats exposed to CUS, we believe this can be extended to all instances where the SPT is used to assess anhedonia. By addressing the confounding effect of metabolic factors in the results of the test, even when food and water are available, researchers increase the robustness of their results and prevent misleading inferences that have detrimental effects in the translatability of the research and its ethical implications.

## Acknowledgements

We would like to thank Dr. Alexandra Bannach-Brown from CAMARADES Berlin for her helpful feedback on the methodology and in the early stages of interpreting the results, Huandi Xu for his help in obtaining manuscripts published in Chinese journals, and all the people that took the time to provide us with more information and raw data from their studies.

