## Supplementary material for "Reliability of sucrose preference testing following short or no food and water deprivation - a Systematic Review and Meta–Analysis of rat models of chronic unpredictable stress"

Jenny P. Berrio^1^, Sara Hestehave^2^, Otto Kalliokoski^1^

### Methods

#### Registration and open access data

Access to the pre-registered protocol can be found here: <https://www.crd.york.ac.uk/prospero/display_record.php?RecordID=259015>

Access to the code, additional files and data sets can be found here: <https://osf.io/wdba5/> Throughout the text you will find references to specific files found in this repository.

##### *Deviations from the pre-registered protocol*

Pairwise *post hoc* analyses of experiments that reported more than one SPT outcome measure in the same cohorts of animals were added to the original protocol for this study. These analyses were performed after confirming that more than 4 studies could be included. An additional sensitivity analysis was performed to check for the robustness of the meta-analyses where the restricted maximum likelihood (REML) estimator of heterogeneity was used instead. We performed these extra analyses to further explore our data and to understand the possible effect of outcome measure on the observed pooled effect. As a *post hoc* analysis, ten percent of the studies that were excluded because of long periods of food and/or water deprivation prior to the sucrose preference test were randomly selected to further explore the influence of fasting through a meta-regression analysis. To facilitate interpretation, a simple funnel plot, instead of a contour-enhanced funnel plot is presented. The funnel plot and Egger’s regression of the experiments reporting preference as the outcome were conducted as a *post hoc* analysis. A funnel plot using the (raw) mean difference against its SE is recommended when assessing for small study effects in meta-analysis of preclinical studies because it is not associated with any artifactual asymmetry^1^.

#### Search strategy

In all databases, the search covered a period from 1987 (first CUS protocol) to June 2021. No other pre-defined limits were used. Each search string containing keywords and indexed terms was piloted and adapted to exclude terms that yielded zero results or significantly decreased the specificity of the search. The final three strings were combined to retrieve the list of potential studies in each database.

The complete search strings were as follows:

##### *Pubmed*

1. “Rats”[Mesh] OR “Rat”[tiab] OR “Rats”[tiab] OR “Rattus norvegicus”[tiab]
2. “chronic unpredictable mild stress”[tiab] OR “chronic unpredictable stress”[tiab] OR “chronic mild stress”[tiab] OR “mild stress”[tiab] OR “Chronic Unavoidable stress”[tiab] OR “Unpredictable Chronic stress” [tiab] OR “chronic variable stress”[tiab] OR “chronic varied stress”[tiab] OR “chronic variate stress” [tiab] OR “chronic stress depression model”[tiab] OR “ Chronic stress model”[tiab]
3. “sucrose preference test”[tiab] OR “sucrose consumption test”[tiab] OR “sucrose intake test” OR “saccharine preference test”[tiab] OR “saccharin preference test”[tiab] OR “sucrose preference”[tiab] OR “saccharine preference”[tiab] OR “saccharin preference”[tiab] OR “sucrose consumption”[tiab] OR “sucrose intake”[tiab] OR “anhedonia”[tiab]

##### *Embase*

1. exp rat/ or “rats”.mp. or “rat”.mp. or “Rattus norvegicus”.mp. or “R. norvegicus”.mp.
2. exp chronic unpredictable stress/ or “CUMS”.mp. or “chronic unpredictable mild stress”.mp. or “chronic mild stress”.mp. or “mild stress”.mp. or “Chronic Unavoidable stress”.mp. or “chronic unpredictable stress”.mp. or “Unpredictable Chronic stress”.mp. or “chronic variable stress”.mp. or “chronic varied stress”.mp. or “chronic variate stress”.mp. or “chronic stress depression model”.mp. or “Chronic stress model”.mp.
3. exp sucrose preference test/ or “sucrose preference test”.mp. or “sucrose consumption test”.mp. or “sucrose intake test”.mp. or “saccharine preference test”.mp. or “saccharin preference test”.mp. or “sucrose preference”.mp. or “saccharine preference”.mp. or “saccharin preference”.mp. or “sucrose consumption”.mp. or “saccharine consumption”.mp. or “saccharin consumption”.mp. or “sucrose intake”.mp. or exp anhedonia/ or “saccharin intake test”.mp or “saccharin intake”.mp or “saccharine intake”.mp

##### *Web of science*

1. TS=(rat$ OR “Rattus norvegicus” OR “R. norvegicus”)
2. TS=(“chronic unpredictable mild stress”) OR TS=(“chronic unpredictable stress”) OR TS=(“chronic mild stress”) OR TS=(“mild stress”) OR TS=(“chronic exposure to stress”) OR TS=(“Unpredictable Chronic stress”) OR TS=(“chronic variable stress”)OR TS=(“chronic varied stress”)OR TS=(“chronic variate stress”) OR TS=(“chronic stress depression model” ) OR TS=(“Chronic stress model”)
3. TS=(“sucrose preference test”) OR TS=(“sucrose consumption test”) OR TS=(“sucrose intake test”) OR TS=(“Saccharin preference test”) OR TS=(“saccharin consumption test”) OR TS=(“Saccharin intake test”) OR TS=(“sucrose preference”) OR TS=(“Saccharin preference”) OR TS=(“Saccharine preference”) OR TS=(“Sweet preference”) OR TS=(“sucrose consumption”) OR TS=(” Saccharin consumption”) OR TS=(“Sucrose intake”) OR TS=(“Saccharin intake”) OR TS=(“anhedonia”)

#### Eligibility criteria

**Supplementary table 1**. *Reasons for exclusion at each stage of screening.*

| **Title and abstract** |  | **Full-text screening** |
| --- | --- | --- |
| **Wrong publication type:**   - Not an original study - Systematic review/Meta-analysis |  | **Wrong publication type** |
|  |  | **Not easily readable** (even with the use of translation tools) |
|  |  | **Insufficient info:**  Report of study did not present sufficient information to assess the fulfillment of all the criteria for inclusion |
| **Wrong study design:**   - Not an animal study |  | **Wrong design:**   - Study did not use a **“control vs exposed”** design - Study uses the exposed group as their own control   **Not appropriate controls:**   - Control differs from the exposed group in more than the exposure variable |
| **Wrong population:**   - Not a study on live laboratory rats - The study does not employ a stress paradigm - The study employs other forms of chronic stress different from CUS |  | **Not adequate rat model:**   - *Wrong population* (mice, pre-weaned rats, ovariectomized females, gonadectomized males, pregnant or post-partum rats, mutant/genetically modified strains) - CUS was not administered exclusively for at least two weeks. - Study at the start of the CUS, co-administers addictive substances or any other drug, compound or special diet - Study administers CUS following another stress protocol or model-inducing intervention AND does not include control- exposed groups in which CUS is exclusively applied |
| **Wrong outcome:**   - Study does not assess behavioral endpoints - Study does not assess anhedonia-like behaviors |  | **Not appropriate intervention:**   - Study does not perform a SPT - Study did not have a familiarization phase to the sweet solution (sucrose, saccharin) |
|  |  | **Long food and water deprivation:**  Study implemented periods of food and/or water deprivation longer than 6 hours before testing |
|  |  | **Wrong SPT timing:** SPT was not performed at the end of CUS or at the end of the antidepressant treatment. |

#### Selection process:

For the numeric account of the selection process, see *(OSF: “Batch summary.xlsx”)*

#### Study details and outcome data extraction:

Supplementary table 2 presents the list of the methodological details and outcomes extracted from each study. A random sample of 25% of the studies was reviewed for extraction accuracy by a second reviewer. Errors were found in only 1.3% of the reviewed data, all in three variables: length of stress, type of test and timing of the test. These three variables were checked again for accuracy for all included studies by one reviewer.

**Supplementary table 2.** *Methodological details and outcomes extracted from included experiments. A shorter list of items, highlighted in blue, were extracted for a subset of studies using longer periods of food and water deprivation.*

| A. **Methodological details** | |
| --- | --- |
| **Study ID** | *Title  *Corresponding author and e-mail address  *Journal  *Year |
| **Subject characteristics** | *Strain  *Sex  *Age at the start of the study*(≤ 9 weeks, ≤ 12 months, > 12 months)  *Weight |
| **CUS Model characteristics** | *Length (in weeks) |
| **SPT characteristics** | * Type of sweet: Sucrose or Saccharine  * Type of test: One-botte, Two bottle  * Length (h) of food and water deprivation, if any.  * Timing: light, dark cycle  * Length of test (h)  * Concentration of sucrose/saccharine (w/v) |
| **Antidepressant treatment (if appropriate)** | *Antidepressant  *Dose (mg/kg)  *Class  *Length of CUS before treatment (in weeks)  *Length of antidepressant treatment (in weeks) |
| B. **Outcome** | |
| **Sweet preference/ consumption at the end of the CUS protocol** | *Stressed vs unstressed control  *Treated vs untreated stressed control  Measuring units:   1. Sucrose/saccharine preference (in percent), or 2. Sucrose/Saccharine intake calculated as weight or volume of sucrose solution (g or mL), or 3. Ratio of sucrose/saccharine intake to total water intake, or 4. Sucrose/Saccharine intake in relation to the animal’s body weight (g/kg), or 5. Other |

(*) When the age was not reported, body weight was used to estimate it based on predefined ranges obtained from growth curves.

Ten percent of the studies that were excluded because they employed periods of fasting longer than 6 hours were randomly selected, and checked against our inclusion and exclusion criteria. A total of 73 studies did not fulfill our criteria and were replaced by randomly selecting another study belonging to the same batch. A total of 73 studies were included in the present investigation and the details colored in blue were extracted by one reviewer. The extracted data was double-checked for extraction accuracy by a second reviewer.

Age was defined as follows:

- **≤ 9 weeks:** P21-P63
- **Adult ≤ 12 months:** P64-P336 (10 weeks to 48 weeks)
- **Adult > 12 months:** > 48 weeks

When the age was not reported, body weight was used to estimate it based on predefined ranges obtained from growth curves from breeders *(OSF: “Weight2Age.xlsx”)*.

The following limits were used for defining adults:

| Strain/stock | Female | Male |
| --- | --- | --- |
| Wistar | ≤ 300g | ≤ 400g |
| Sprague-Dawley | ≤ 250g | ≤ 350g |
| Long-Evans | ≤ 250g | ≤ 350g |
| Lister-Hooded | ≤ 200g | ≤ 300g |

In cases where the age or weight was reported as a range, the upper limit was used to select the developmental stage.

Supplementary table 3 presents the considerations agreed upon and followed when performing the outcome data extraction in relation to different study designs and data reporting. When outcome data was extracted from figures, the average of the two values obtained by the reviewers was used in the final data set. After contacting the authors in cases of missing data, a waiting period of three weeks was established before deciding to exclude the study. In no instances did we receive replies past this point.

**Supplementary table 3**. *Outcome extraction considerations regarding specific data reporting cases and study designs.* These considerations and actions were agreed upon by the researchers prior to starting the dual extraction.

| **Data reporting** | | **Study design** | |
| --- | --- | --- | --- |
| **Consideration** | **Action** | **Consideration** | **Action** |
| a) n reported in ranges | Lowest value was extracted | a) Studies with re-exposure to CUS | Data was extracted from the first test performed after the end of the first CUS |
| b) SD is not reported | Standard error of the mean (SEM) or confidence intervals were used to calculate it | b) Studies where a secondary intervention was started during the CUS and no placebo/sham group was used | Data was extracted from the last test performed before the start of the new intervention |
| c) It is unclear what the error bars represent in figures | SEM was assumed (for conservative estimate) | c) Studies with several control and/or stress groups AND where one-to-one control vs stress comparisons are not straightforward | Data were combined to obtain an aggregate measure for one-to one comparison  Exception: whenever groups vary in variables of interest (see data analysis), the data were extracted for each group in reference to a single control or stress group and the number of animals was adjusted in the shared group by the number of comparisons^2^. |
| d) Sucrose and saccharine data for the same cohorts of animals are reported | Sucrose data was extracted as the main outcome and Saccharine as the alternative outcome | d) Studies with repeated measurements of SPT | Data extracted from the first test performed after the end of the CUS |
| e) Study reports different SPT units | Extract measuring unit by priority:  1. Sweet preference (%)  2. Sweet intake/weight (g or mL/kg)  3. Sweet intake (g or mL)  4. Ratio of sweet intake to total water intake  5.Other  Whenever 2 and/or 3 were reported in addition to 1, they were extracted as alternative outcomes. | e) Study reports different sweet concentrations | Data for concentrations between 1-2% were extracted. If these concentrations were not used, the data of the concentration tested closest to the end of the CUS was selected |
| f) Study reports data for “stress resilient” and “stress susceptible” animals separately | Data were combined to obtain an aggregate measure for the stress group |  |  |

#### Quality and risk of bias assessment:

##### *Quality of reporting*

**Supplementary Table 4**. *Quality of reporting checklist.*

| Quality reporting (modified from ARRIVE essential 10) | |
| --- | --- |
| Item | Description |
| **Experimental animals** | The study provides details on the strain/stock, sex, age/weight at the beginning of the experiment, and the provenance of the experimental subjects. |
| **Study design** | The study clearly states the experimental unit and describes all groups being compared. |
| **Sample size** | The study reports the exact number of experimental units allocated to each group at the start of the experiment AND explains how the sample size was decided on (*e.g.,* *a priori* sample size calculation). |
| **Allocation** | The study clearly states the strategy used to assign experimental units to the groups (*e.g.* randomization or matching) |
| **Blinding** | The study reports performing blinding at any stage of the experiment. |
| **Definition of Outcome** | The study clearly states what the outcome measure of the sucrose preference test was, and how it was obtained, in the text or graphs. |
| **Statistical methods** | The study makes it clear which statistical method was used to compare the sucrose preference test across groups, including *post hoc* comparison methods (if used). |
| **Experimental procedures** | Both of the following need to be present to consider this item to be adequately reported:   a. **Stress protocol:** The study describes the type of stressors used in the chronic unpredictable stress protocol and their general schedule. b. **Sucrose test:** The study describes how pre-exposure to sucrose/saccharine was performed (if done) and the study describes the sucrose preference test, its frequency and timing. |
| **Exclusions** | The study clearly indicates within the text or in figures, the exact number of experimental units analyzed for the sucrose preference test. If this number does not match the initial sample, the study explains why the experimental units were not included in the analysis. |
| **Results** | Data of the SPT with descriptive statistics (group sizes, measures of central tendency and a measure of variability) for all groups is reported within the text, in tables or graphs, or in a supplementary file. |

Access to the data: *(OSF: “QR_Final.xlsx”)*

##### *Risk of bias*

**Supplementary Table 5**. *Risk of Bias checklist.* Attrition and reporting biases were assessed as single items.

| Risk of bias (modified from SYRCLE’s risk of bias tool for animal studies) | | |
| --- | --- | --- |
| Bias | Item | Signaling questions |
| Selection | **Randomization** | Did the investigators describe a random component in the allocation of experimental units? |
|  | Sequence | Did the investigators use a clear strategy to allocate the experimental units? |
|  | **Allocation concealment** | Was the allocation sequence (the group each individual animal is to be allocated to) adequately concealed from the experimenter prior to the allocation? |
|  | Baseline Characteristics | Was the distribution of sex, weight, sucrose preference (Baseline SPT) balanced for the intervention and control groups? (Either matched at the beginning or reported in the results). |
| Performance | **Stress exposure** | Were possible confounding factors related to stress exposure accounted for between the control and exposed groups? |
|  | **Blinding** | Studies with antidepressant treatment only: Were the investigators blinded to which intervention (placebo vs active treatment) each animal received during the experiment? |
| Detection | **Random outcome assessment** | If applicable (studies employing a one-at-a-time testing), did the investigators test the animals in a random order during the outcome assessment? |
|  | **Place preference** | Were the bottles counterbalanced during the test to avoid place preference? |
|  | **Blinding** | Was blinding of the outcome assessor ensured? |
| Attrition | **Attrition** | If there are any missing data, are they adequately explained and dealt with transparently? |
| Reporting | **Reporting** | Does the study report the SPT outcome with descriptive statistics (group sizes, measures of central tendency and a measure of variability) for all groups compared? |
| **Scoring system:** Each item was assessed as Yes: explicitly explained in the paper; No: explicitly said or easily deduced from the report that it was not done; Unclear: Not explained and it is hard to judge if this was done or not. The sum of all items answered with a “Yes” was considered the risk of bias score. A higher number indicates more actions taken to reduce bias. Performance blinding was only assessed for experiments employing an antidepressant treatment. Consequently, the risk of bias score sums to a maximum of 10 in experiments comparing stressed versus control groups, and a maximum of 11 in experiments comparing stressed versus treated groups. | | |

#### Data synthesis

The following R packages were used in the data analysis and the graphics:

1. Package *tidyverse*: Wickham H, Averick M, Bryan J, Chang W, McGowan LD, François R, Grolemund G, Hayes A, Henry L, Hester J, Kuhn M, Pedersen TL, Miller E, Bache SM, Müller K, Ooms J, Robinson D, Seidel DP, Spinu V, Takahashi K, Vaughan D, Wilke C, Woo K, Yutani H (2019). “Welcome to the *tidyverse*.” *Journal of Open Source Software*, *4*(43), 1686. <doi:10.21105/joss.01686> <https://doi.org/10.21105/joss.01686>.
2. Package *meta*: Balduzzi S, Rücker G, Schwarzer G (2019), How to perform a meta-analysis with R: a practical tutorial, Evidence-Based Mental Health; 22: 153-160.
3. Ushey K (2022). *renv: Project Environments*. R package version 0.15.5, <https://CRAN.R-project.org/package=renv>.
4. Harrer, M., Cuijpers, P., Furukawa, T. & Ebert, D. D. (2019). dmetar: Companion R Package For The Guide ‘Doing Meta-Analysis in R’. R package version 0.0.9000. URL <http://dmetar.protectlab.org/>.
5. Wickham H, Bryan J (2022). *readxl: Read Excel Files*. R package version 1.4.0, <https://CRAN.R-project.org/package=readxl>.
6. H. Wickham. ggplot2: Elegant Graphics for Data Analysis. Springer-Verlag New York, 2016.
7. Xiao N (2018). *ggsci: Scientific Journal and Sci-Fi Themed Color Palettes for ‘ggplot2’*. R package version 2.9, <https://CRAN.R-project.org/package=ggsci>.
8. Kassambara A (2020). *ggpubr: ‘ggplot2’ Based Publication Ready Plots*. R package version 0.4.0, <https://CRAN.R-project.org/package=ggpubr>.
9. Zhu H (2021). *kableExtra: Construct Complex Table with ‘kable’ and Pipe Syntax*. R package version 1.3.4, <https://CRAN.R-project.org/package=kableExtra>.
10. Neuwirth E (2022). *RColorBrewer: ColorBrewer Palettes*. R package version 1.1-3, <https://CRAN.R-project.org/package=RColorBrewer>.
11. Slowikowski K (2021). *ggrepel: Automatically Position Non-Overlapping Text Labels with ‘ggplot2’*. R package version 0.9.1, <https://CRAN.R-project.org/package=ggrepel>.
12. Wilke C (2020). *cowplot: Streamlined Plot Theme and Plot Annotations for ‘ggplot2’*. R package version 1.1.1, <https://CRAN.R-project.org/package=cowplot>.
13. Gordon M, Lumley T (2021). *forestplot: Advanced Forest Plot Using ‘grid’ Graphics*. R package version 2.0.1, <https://CRAN.R-project.org/package=forestplot>.

##### *Power calculation*

Power calculations were performed with the function power.analysis in the R package *dmetar*^3^

A meta-analysis of ten studies, with mean sample size of 10, have 84% power to detect a standardized effect size (SMD) of 0.6 (alpha level: 0.05), in a random-effects model, if high (I^2^ > 75%) heterogeneity is assumed.

The power of a meta-analysis of 4 studies per subgroup, with a mean sample size of 10, to detect a standardized effect size of 1.0 in a random-effects model (using an alpha-level of 0.05 and assuming high heterogeneity) is 85%. Considering that a previous review of the SPT found an standardized mean difference between the stress and control groups in sucrose consumption/intake of no less than 1.5 SMD in multiple subgroup comparisons^4^, 4 studies per subgroup were considered to the minimum requirement for performing an analysis.

##### *Data aggregation*

If a study included several control or stress groups and no “one-to-one” control vs stress comparison was possible (*e.g.*, one control group is compared to two stressed groups that fulfilled our criteria), the groups were combined using the recommendations in the Cochrane Handbook^2^ to obtain an aggregate measure (*i.e.*, the stressed groups were combined to obtain a single measure that was compared to the control group). Only in one study, the groups were not combined because the groups varied in a variable of interest – the length of the stress protocol (Supplementary table 2, Study design (c)). In this case, the ‘shared control’ group was split into two to avoid double-counting^2^.

One study reported the outcomes as medians with interquartile ranges. In this case the median was assumed to approximate the mean, and the SD was calculated from the IQR^2^.

Different measures of consumption per body weight were converted into g/kg.

##### *Pairwise analyses:*

Pairwise analyses were conducted for experiments that reported preference and intake or preference and intake corrected by body weight in the same cohorts of animals. For these analyses, the pooled effect of the difference in effect sizes between stressed animals and controls for both outcome measures was estimated using a fixed effects model. The sample size for both outcomes was calculated as the sum of the sample size in the stressed and control groups divided by two.

#### Sensitivity analyses:

Two sensitivity analyses were performed:

###### Sample size:

When the sample size of the studies was reported in ranges, the lowest value was extracted for the primary analysis (Supplementary table 2, Data reporting (a)). For sensitivity, the analysis was re-run using the highest number instead (25 experiments).

###### Heterogeneity estimator:

The restricted maximum likelihood (REML) estimator was recently recommended over the DerSimonian and Laird (DL) estimator for gauging between-study heterogeneity in meta-analyses of preclinical data^5^. To test the robustness of our results, we re-ran the primary analysis using this estimator instead.

### Results

**Databases:** *(OSF: “Database_Main” & “Database_alt”)*

**List of references of included studies:** *(OSF: “Bibliography_Included studies”)*

##### *Additional characteristics of included experiments*

**Characteristics of the experiments included in the analysis**

See *OSF: “Tables.html”*

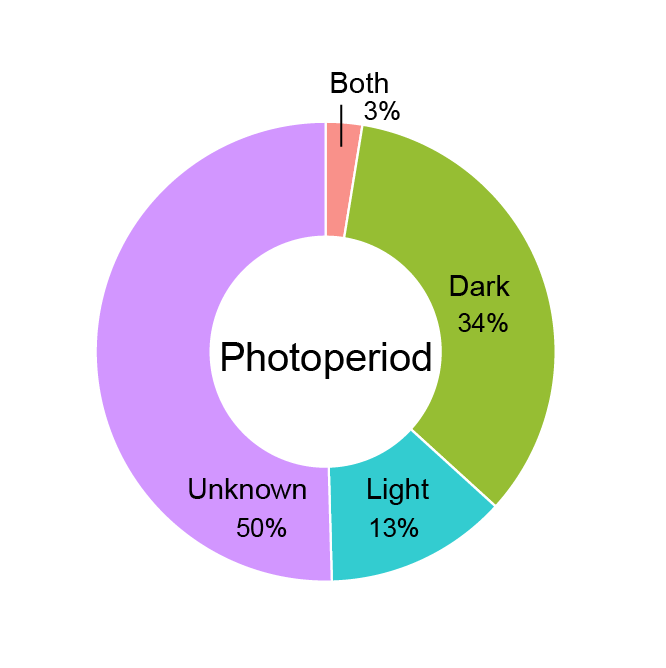

**Supplementary figure 1.** *Distribution of the experiments with tests shorter than 12 h by photoperiod*. 34% (40 out of 117) performed the test during the dark phase, 13% (15) during the light phase, and 50% (59) did not report the photoperiod. Only 3 experiments (1 study) had the test spread over both phases.

##### *Quality of reporting*

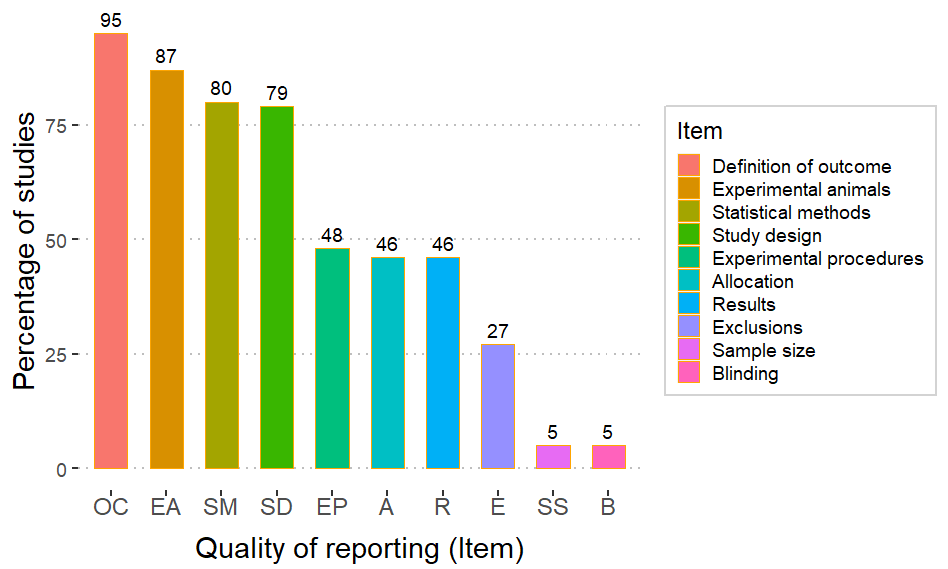

**Supplementary figure 2.** *Quality of reporting*. Proportion of studies that reported the individual items assessed in the quality of reporting checklist adapted from the ARRIVE essential 10. Each item is abbreviated on the X-axis. *(OSF: “QR_Final.xlsx”)*

##### *Risk of bias*

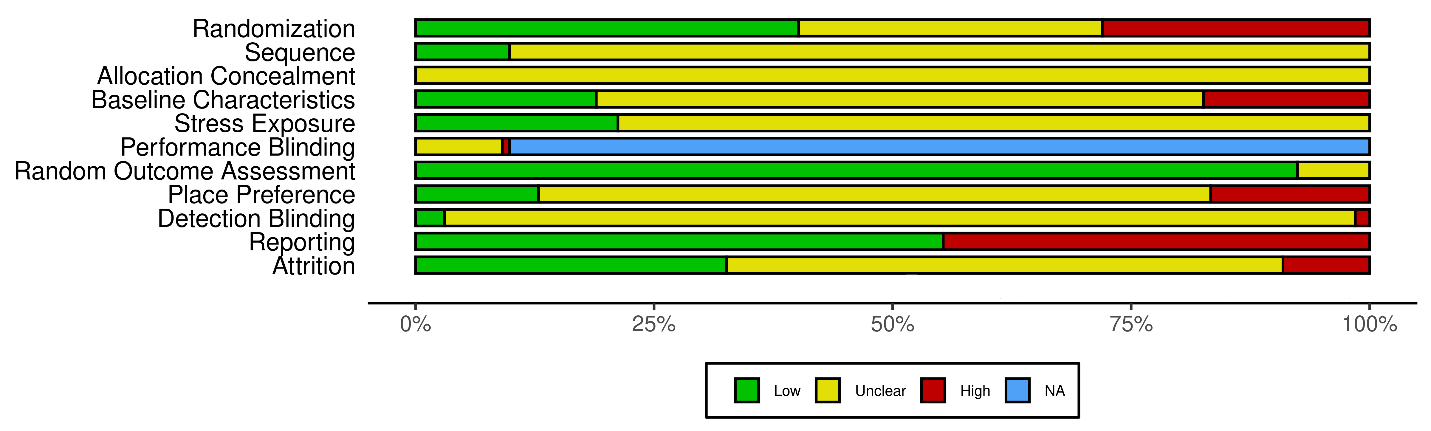
**Supplementary figure 3.** *Risk of bias*. Assessed using a modified version of SYRCLE’s risk of bias tool. The proportions of the colored bars indicate the percentage of studies (n = 132) with low (green), unclear (yellow) and high (red) risk of bias for each item. NA stands for “not applicable”. Only the studies that used antidepressant treatment were evaluated for performance blinding (n = 12). The shiny web app Robvis was used to create the risk of bias plot^6^.

###### Selection bias:

40% of studies (53 studies) mentioned a random component to the allocation of the experimental units, while only 9.8% (13) explained the allocation strategy. None of the studies reported allocation concealment. The sex (either only one sex was employed or experimental cohorts were balanced with respect to the subjects’ sex when both sexes were included in the same experiment), weight, and baseline sweet preference of the subjects across groups were balanced in 18.9% (25), while they were not in 17.4% (23). For most studies, it was unclear if the groups were similar in these key characteristics at baseline (63.6%/84 studies).

###### Performance bias:

21.2% of studies (28) housed the controls and stress groups separately but under similar environmental conditions to avoid indirect stress exposure of controls. No studies included in the analysis of antidepressant treatments mentioned blinding with respect to the therapeutic intervention, except for one study that clearly stated that blinding was not used.

###### Detection bias:

In most studies, the SPT was carried out in the home cage, and the subjects were assessed simultaneously. Less than 1% of the studies (10) had a testing methodology that could introduce an effect of testing order earning them an “unclear risk” in the random outcome assessment item. 15.9% of studies (21) properly counterbalanced the bottles within the test period to avoid place preference. Studies that performed the counterbalancing outside of the testing period or employed a one-bottle paradigm were classified as high risk (16.6%/22). The outcome assessors were blinded in only 4 studies.

###### Reporting and attrition bias:

55.3% of studies (73) reported the SPT outcome with descriptive statistics and variability for all groups. 32.6% of studies (43) had no missing data or explained any exclusions clearly. In 58.3% of studies (77) the lack of reporting of the initial and final sample sizes rendered the assessment of attrition bias unclear. The remaining 12 studies (9.1%) had unexplained missing data (high risk).

Access to the data:

1. Robvis scale: *(OSF: “RB_Final_ROBVIS.xlsx”)*
2. RoB score: *(OSF: “RB_Final_Score.xlsx”)*

##### *Data synthesis*

**Full forest plot**

Pdf of forest plot: *(OSF: “forestplot_Q1_all”)*

**Supplementary figure 4.** *Forest plot of experiments comparing the sweet intake/preference in stressed animals to unstressed controls*. The names of the experiments are composed of the main author of the study, year of publication and an indicator of the number of experiments extracted from a single published study (“_Cx”).

**Pairwise analyses**

Access to the data:

1. *Preference and intake for the same animals*: *(OSF: “Preference_pairwise.xlsx” & “Intake_pairwise.xlsx”)*
2. *Preference and intake corrected by body weight for the same animals*: *(OSF: “Pref_weight_pairwise.xlsx” & “corrected_weight_pairwise.xlsx”)*

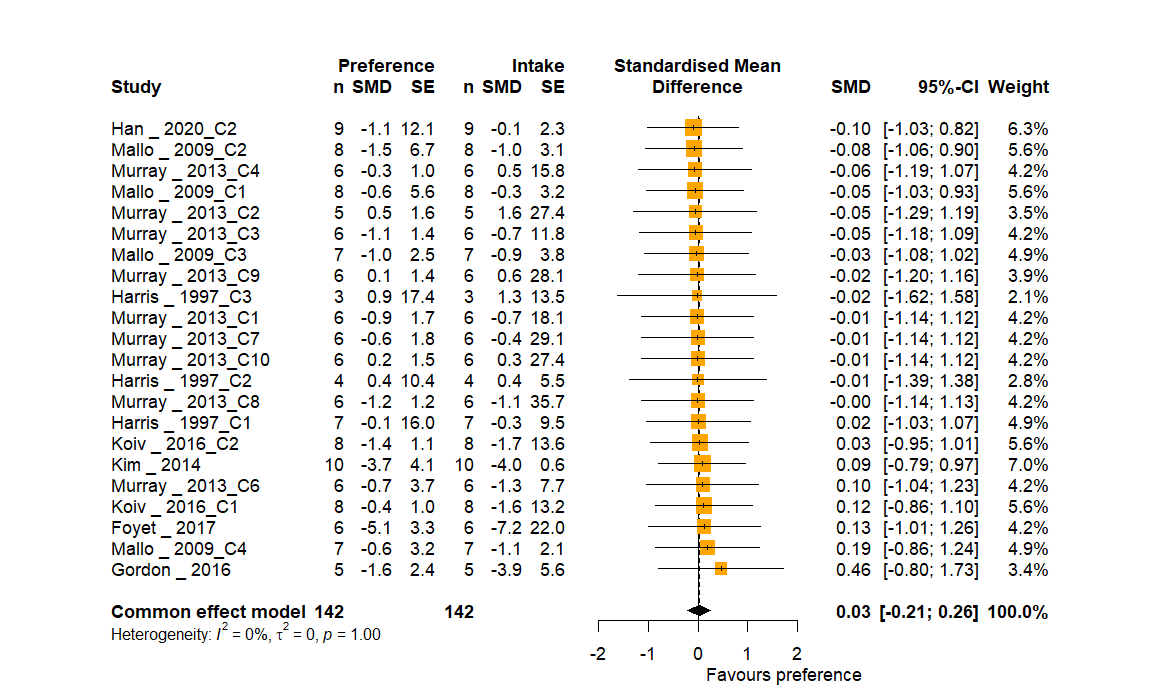

**Supplementary figure 5.** *Forest plot of experiments that reported preference and intake for the same animals*. The differences in effect sizes between the outcome measures were pooled with a fixed effects model. The pooled effect and its 95% CI is represented by the black diamond. Abbreviations: n, number of animals in each group; SMD, standardized mean difference; SE, Standard error; 95%-CI, 95% confidence interval.

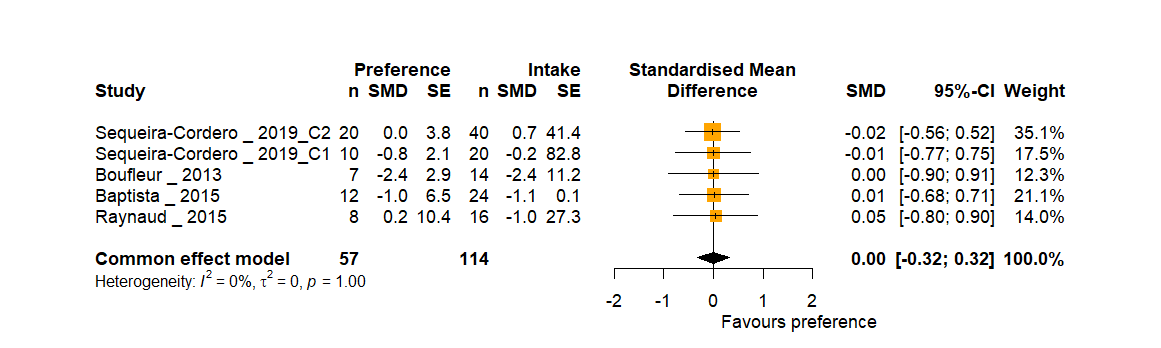

**Supplementary figure 6.** *Forest plot of experiments that reported preference and intake corrected by body weight for the same animals.* The differences in effect sizes between the outcome measures were pooled with a fixed effects model. The pooled effect and its 95% CI is represented by the black diamond. Abbreviations: n, number of animals in each group; SMD, standardized mean difference; SE, Standard error; 95%-CI, 95% confidence interval.

**Other sources of variability**

**Supplementary table 6.** *Summary univariate meta-regression analysis*.

| **Variable** | **Coefficient (SMD)** | **95% CI** | ***z*** | **p value** | **R^2^** | **I^2^** |
| --- | --- | --- | --- | --- | --- | --- |
| **Stressed vs Control** | | | | | | |
| Length of stress (weeks) | -0.125 | [-0.24, -0.009] | -2.1 | 0.035 | 1.67 % | 78.83% |
| Length of food and water deprivation (hours) (studies using period of fasting ≤ 6 h) | -0.026 | [-0.11, 0.06] | -0.63 | 0.53 | 0.47 % | 79.05% |
| Length of food and water deprivation (hours)  (**+ studies with fasting > 6 h**)  Accounting for type of sweetener and length of the stress protocol | -0.029 | [-0.05, - 0.01] | -3.62 | 0.0003 | 3.16% | 80.37% |
|  | -0.027 | [-0.04, - 0.01] | -3.25 | 0.0012 | 2.32% | 79.77% |
| Risk of bias score | 0.021 | [-0.09, 0.14] | 0.36 | 0.72 | 0 % | 79.27% |
| **Treated vs stressed** | | | | | | |
| Length of antidepressant treatment (weeks) | 0.044 | [-0.63, 0.72] | 0.13 | 0.9 | 0 % | 76.67% |
| Length of stress (weeks) | -0.119 | [-0.82, 0.58] | -0.33 | 0.73 | 0 % | 76.91% |
| Length of food and water deprivation (hours)  **(only** **studies with fasting ≤ 6 h)** | -0.21 | [-0.63, 0.20] | -0.99 | 0.32 | 0 % | 76.36% |
| Risk of bias score | -0.26 | [-1.09, 0.57] | -0.61 | 0.54 | 0 % | 76.62% |

R^2^: the amount of heterogeneity accounted for by the predictor. I^2^: the variance not explained by the predictor (remaining between-study heterogeneity). Abbreviations: 95% CI, 95% confidence interval. Stressed vs Control: 171 experiments; treated vs stressed: 12 experiments. Abbreviations: SMD, Standardized mean difference.

Access to the data used to perform the meta-regression for duration of food and water deprivation on the effect measured in the SPT:

*OSF: “DataQ1_plusLFWd.xlsx”*

###### Characteristics of the studies with fasting exceeding 6 hours

From the studies excluded because of long food and/or water deprivation (728, figure 2), 73 reports were randomly selected. A total of 89 experiments were extracted. Wistar and Sprague Dawley rats were the most common stocks in the experiments (40.4% and 53.9%, respectively). Only five experiments used other strain/stocks (Fischer 344, Lister hooded, and Wistar-Han). Male rats were employed in 93.2% of the experiments; only five experiments used females. The length of stress varied from 2 to 11 weeks (51.7% between 2 and 4 weeks, 20.2% between 4 and 6, 21.34% between 6 and 8, and 6.7% above 8 weeks). All experiments used sucrose as sweetener and employed periods of fasting that ranged from 10 to 24 hours. Most (66.3%) deprived the animals of food and/or water for 20 hours or more, 9% between 15 and 19 hours and 24.7% between 10 and 14 hours. The allotted time for consumption in the vast majority of the experiments (70.8%) was half to one hour. Only 10 experiments employed prolonged periods of testing (12 to 48 hours). Overall, with the exception of the expected differences in length of fasting and testing, and a slight skewness towards employing longer periods of stress, these experiments have similar model and methodological characteristics as those employing shorter deprivations.

*Multiple meta-regression:*

To assess whether the type of sweet or the duration of the stress could have confounded the results of the meta-regression. We removed the experiments that have employed saccharin and conducted a multiple meta-regression controlling for length of stress. Length of food and water deprivation was indeed associated with higher effects (decreased sweet consumptions)(Supplementary table 6). Duration of stress was not a significant predictor (-0.0474 SMD, 95%CI [-0.14, 0.04], p = 0.29). This means that the relationship is not confounded by the fact that studies employing longer periods of deprivation may have employed longer periods of stress.

**Sucrose preference test in rats treated with antidepressants:**

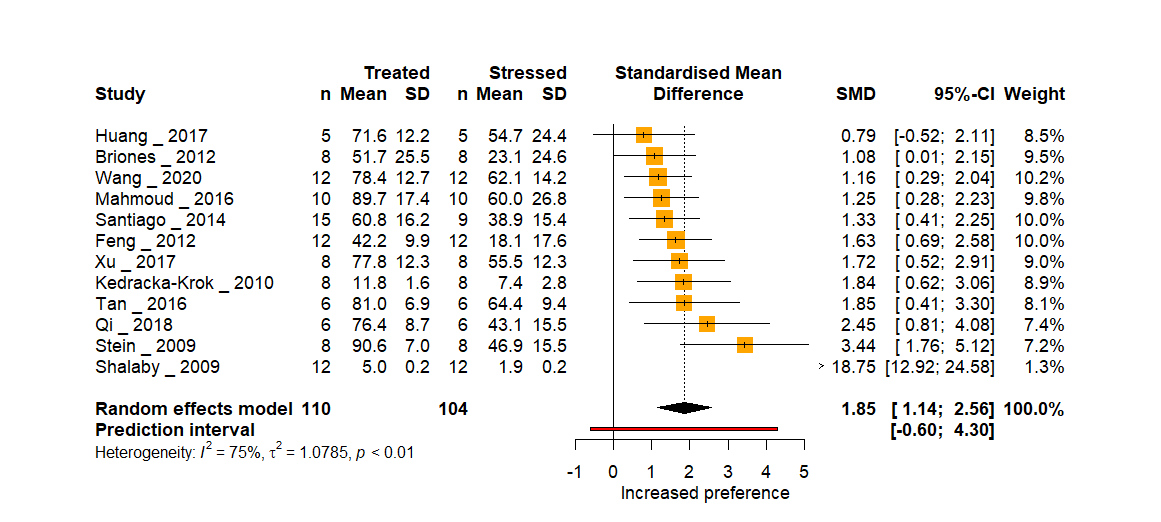

**Supplementary figure 7.** *Forest plot of the sweet consumption in stressed animals compared to stressed animals treated with an antidepressant.* The pooled effect and its 95% CI is represented by the black diamond. Studies falling on the right side of the no effect line found an increased consumption in treated animals. Abbreviations: n, number of animals in each group; SD, standard deviation; SMD, standardized mean difference; 95%-CI, 95% confidence interval.

**Strain - Treated rats vs stressed rats**

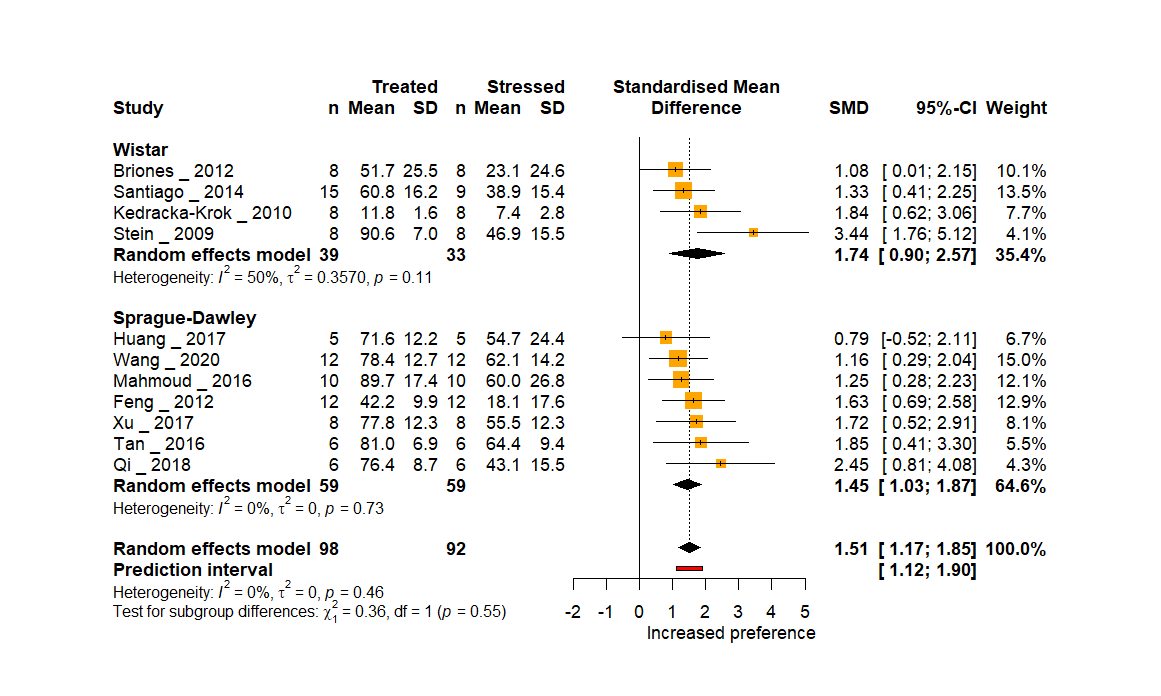

**Supplementary figure 8.** *Subgroup analysis by strain*. The pooled effect in each subgroup and its 95% CI is represented by the black diamond. Studies falling on the right side of the no effect line found an increased consumption in treated animals. Abbreviations: n, number of animals in each group; SD, standard deviation; SMD, standardized mean difference; 95%-CI, 95% confidence interval.

**Class of antidepressant - Treated rats vs stress controls**

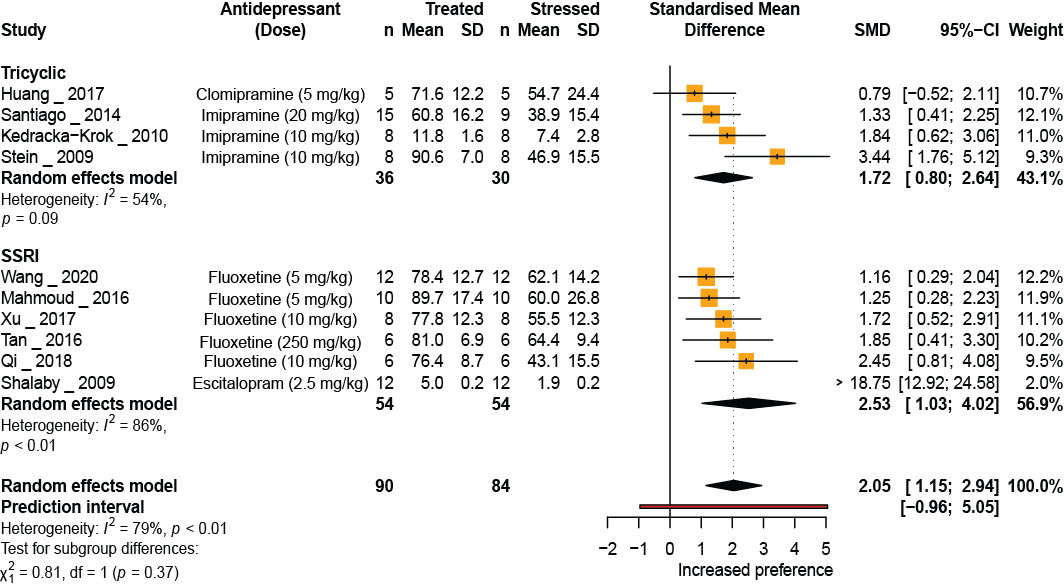

**Supplementary figure 9.** *Subgroup analysis by class of antidepressant*. The pooled effect in each subgroup and its 95% CI is represented by the black diamond. Studies falling on the right side of the no effect line found an increased consumption in treated animals. Abbreviations: n, number of animals in each group; SD, standard deviation; SMD, standardized mean difference; 95%-CI, 95% confidence interval.

##### *Publication bias - small study effects*

The funnel plot is a visual representation of the relationship between the precision (standard error) and the effect size (SMD or MD) of experiments. Experiments with higher precision (smaller standard error) will ideally present effect sizes that cluster around the overall effect calculated. As the precision falls (increasing standard error) the experiments will scatter more widely on either side (symmetrically) as they become worse at estimating the true effect; hence the funnel shape. Small studies producing small effects, null effects, or effects in the opposite direction of the tested hypothesis are, historically, less likely to be published. Researchers obtaining these disheartening results may also choose to add subjects to their study – producing a larger study, with results likely to be closer to the true effect. Regardless of mechanism, these biases, affecting small studies disproportionately, produce funnel plots that are asymmetrical at their base. This is often referred to as the small study effect^7^.

The influence of small studies was initially assessed by visual inspection of funnel plots of the SMD followed by Egger’s test with the corrected standard error (SE) proposed by Pustejovsky and Rogers^8^. In our investigation, the included experiments did not produce a funnel shape and the overall pattern looked asymmetrical (Supplementary figure 10). Experiments with very high effect sizes in the bottom-left corner of the plot are not offset by similarly sized studies in the bottom-right corner, producing, instead, a lopsided funnel shape. What is more, the studies with greater precision in our sample have smaller effect sizes than expected. However, Egger’s regression test with the Pustejovsky correction could not substantiate this assessment (intercept: -1.08, t: -0.21, p = 0.83). As a *post hoc* analysis, we assessed the small study effect only in experiments that reported preference. In this case the mean difference (MD) and its SE were calculated and plotted in the funnel plot. Egger’s test was used to corroborate the results. The asymmetrical pattern in the funnel plot was seen regardless of the effect size metric used (SMD for all included experiments, or the un-standardized mean difference (MD) for experiments reporting the outcome of the SPT as a preference), but the asymmetry was only confirmed by the Egger’s test in the latter case (main manuscript). The Pustejovsky correction is used to prevent the inflation of false positives of asymmetry due to the artifactual correlation between the SMD and its standard error^8^. The correction comes at the cost of reducing overall variation in the precision metric, however. This may be problematic with preclinical studies, where most studies are already fairly small and variation in precision is, consequently, low. Such a method is however not necessary when raw MDs are used. This measure is not associated with any artificial distortion in the funnel plot and therefore it is safe to use in the analysis of small effect study, particularly in meta-analysis of preclinical studies^1^.

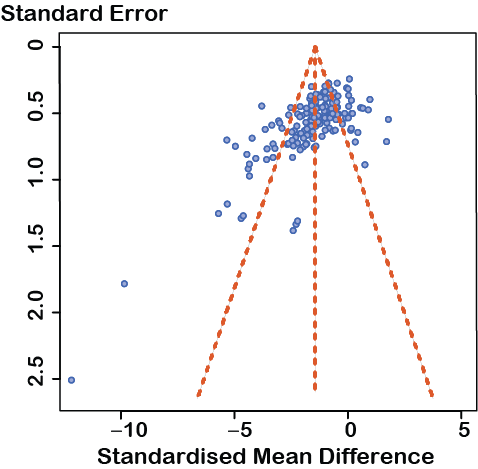

**Supplementary figure 10.** *Funnel plot of experiments including all outcome measures (n = 171) comparing stressed rats to unstressed controls*. The observed effect sizes (SMD) of the experiments (x-axis) is plotted against their standard error (y-axis). The y-axis is inverted with studies with the lowest standard errors on top. This plot, nor those in subsequent funnel plots of the SMD, do not use the standard error proposed by Pustejovsky and Rogers.

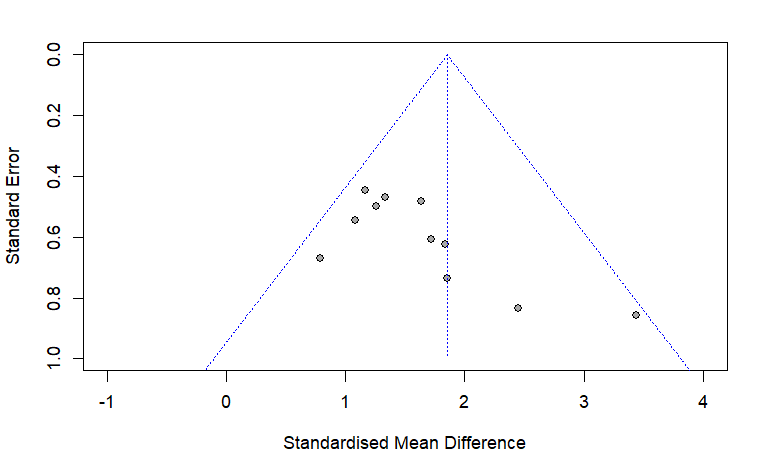

**Supplementary figure 11.** *Funnel plot of experiments comparing antidepressant-treated rats to stressed controls*. The observed effect size (SMD) of the experiments (x-axis) is plotted against their standard error (y-axis). Funnel plot of MD was not carried out because only 9 experiments reported the outcome as a preference. Egger’s regression: intercept: 1.44, t: 0.05, p = 0.96.

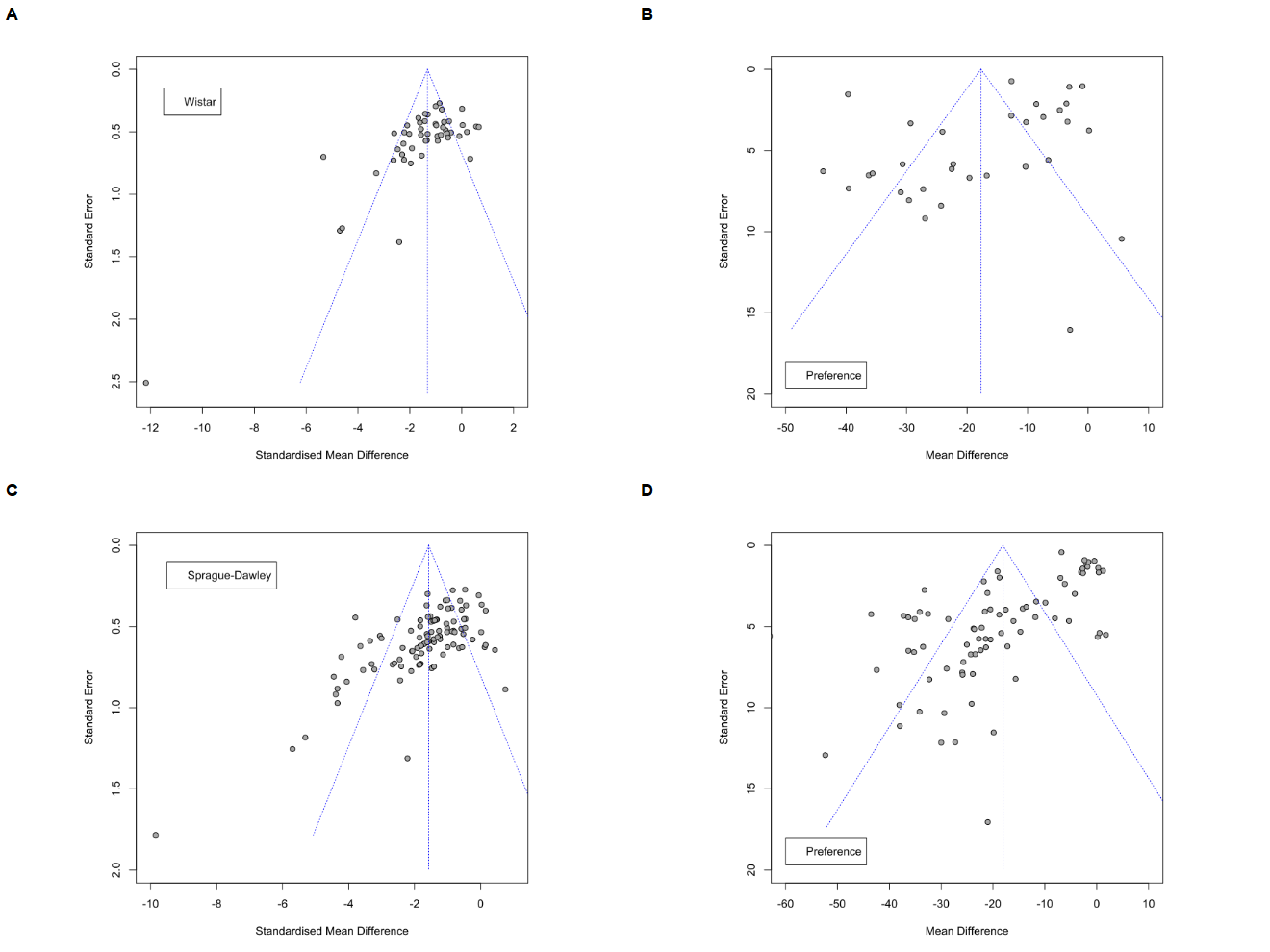

**Supplementary figure 12.** *Funnel plots of experiments comparing stressed rats to unstressed controls stratified by strain.* **A & C** The SMD of experiments (x-axis) using Wistar or Sprague-Dawley rats, respectively, is plotted against their standard error (y-axis). **B & D** The MD of experiments (x-axis) using Wistar or Sprague-Dawley rats reporting their results as a preference is plotted against their standard error (y-axis). In all graphs, the y-axis is inverted so “higher” values represent smaller standard errors.

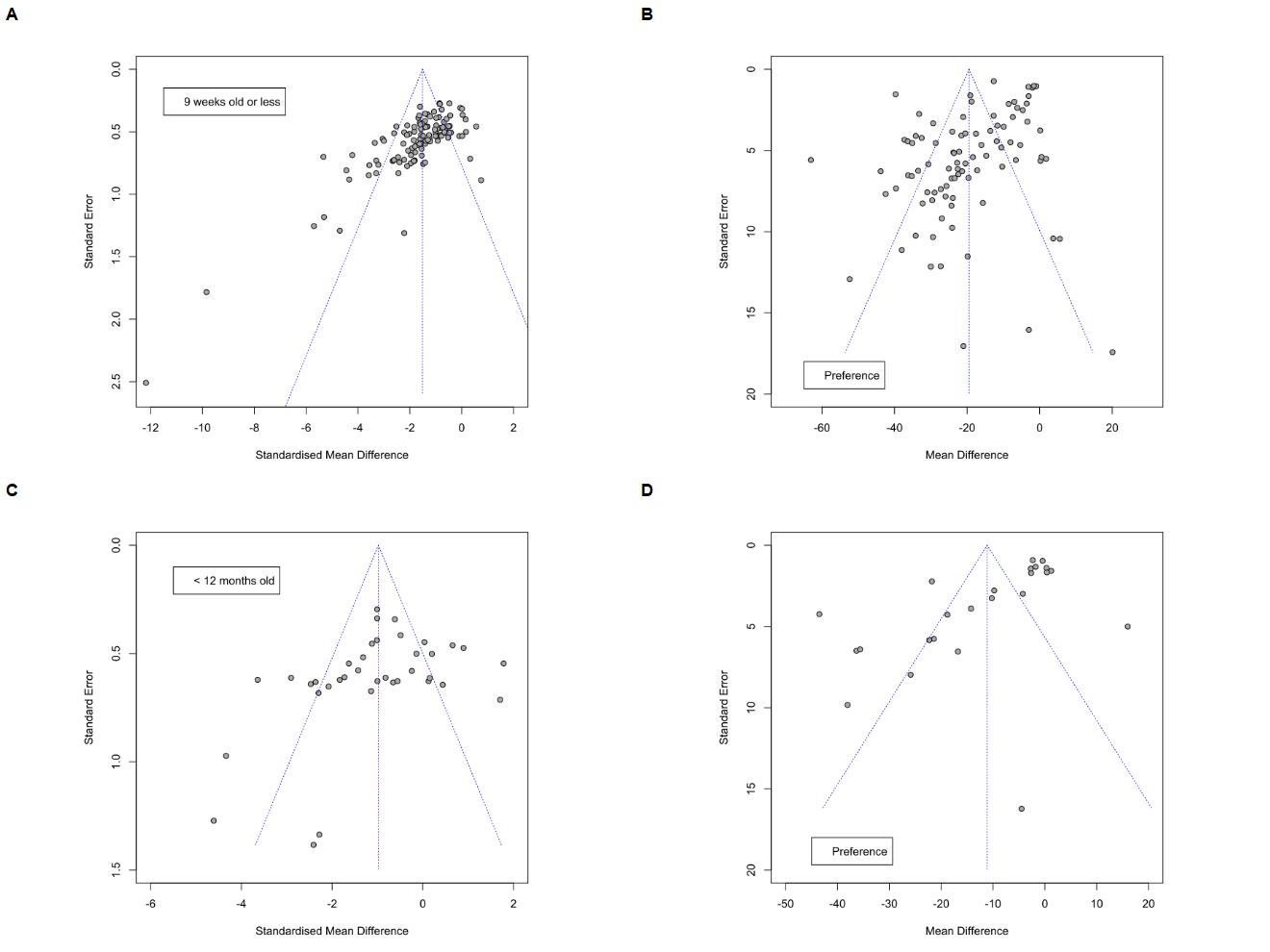

**Supplementary figure 13.** *Funnel plots of experiments comparing stressed rats to unstressed controls stratified by age.* **A & C** The SMD of experiments (x-axis) using rats 9 weeks old or less and adult rats (older than 9 weeks) up to 12 months old, respectively, is plotted against their standard error (y-axis). **B & D** The MD of experiments (x-axis) using rats 9 weeks old or less and adult rats (older than 9 weeks) up to 12 months old that report their results as a preference is plotted against their standard error (y-axis).

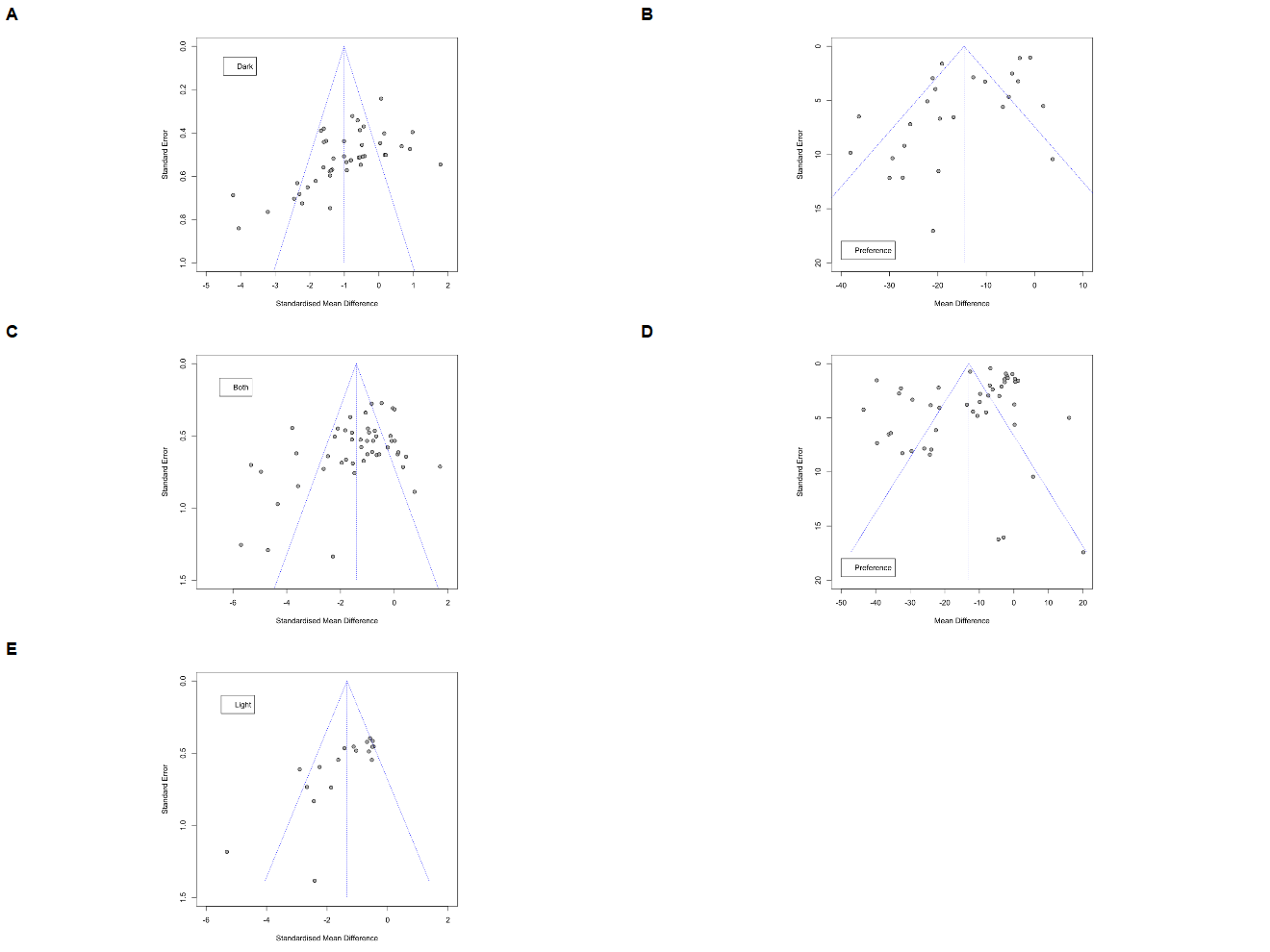

**Supplementary figure 14.** *Funnel plots of experiments comparing stressed rats to unstressed controls stratified by photoperiod of the test.* The SMD of experiments (x-axis) performing the test during the dark period (**A**), light period (**E**), or, both (**C**) is plotted against their standard error (y-axis). The MD of experiments performing the test during the dark period (**B**) or throughout both periods (**D**) that report their results as a preference (x-axis) is plotted against their standard error (y-axis). A funnel plot with MDs for the light period was not carried out because only 6 experiments reported their outcome as a preference.

**Supplementary table 7**. Results of the Egger’s regression test with Pustejovsky correction (_p) by subgroup when SMD was used, and of Egger’s regression test when MD was used for experiments reporting results as preferences.

| Subgroup | Intercept_p | t_p | p_value_p | Intercept | t | p_value |
| --- | --- | --- | --- | --- | --- | --- |
| Wistar | -1.26 | 0.23 | 0.82 | -7.74 | -1.70 | 0.0986 |
| Sprague-Dawley | -1.37 | 0.12 | 0.90 | -2.09 | -6.98 | < 0.0001 |
| < 10 weeks | -1.00 | -0.65 | 0.51 | -5.76 | -4.71 | < 0.0001 |
| < 12 months | -1.41 | 0.73 | 0.47 | 3.20 | -4.01 | 0.0006 |
| light | 0.87 | -1.29 | 0.21 | NA | NA | NA |
| Dark | 0.12 | -1.12 | 0.27 | -2.27 | -3.24 | 0.0037 |
| Both | -1.87 | 1.25 | 0.22 | -5.10 | -1.92 | 0.0620 |

##### *Sensitivity analysis*

###### Sample size

When the sample sizes of the experiments were reported in ranges, the lowest value was used for the primary analyses. The analyses were re-run using the largest possible sample size as well (25 experiments). No significant differences in the results were seen (Supplementary figure 15).

Data: *(OSF: “Database_Main_S” & “Database_alt_S”)*

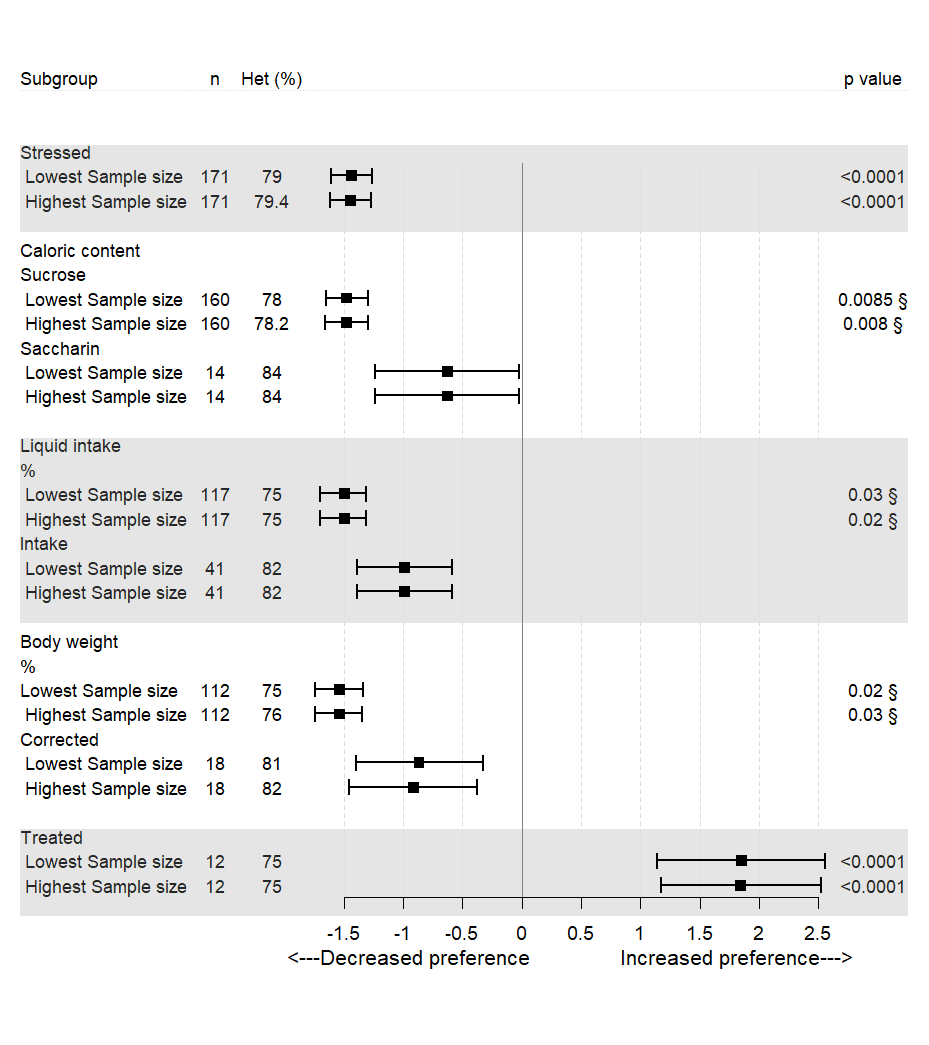

**Supplementary figure 15.** *Forest plot of sensitivity analyses*. The results obtained in each of the analyses performed when the lowest sample size reported was used are shown in comparison to the results of the analyses when the highest sample size was used instead. Similar results were seen in both conditions. The n represents the number of experiments included in each subgroup. § p value of the test for subgroup differences. Abbreviations: Het, between-study heterogeneity (I^2^); %, percentage of sweet preference.

###### Heterogeneity estimator

The restricted maximum likelihood (REML) estimator has been recommended over the DerSimonian-Laird estimator (DL) for assessing between-study heterogeneity in meta-analyses of preclinical data^5^. To test the robustness of our results, we re-ran the primary analyses using this estimator instead. Results were similar to those obtained with the DL method. Stressed rats had a significantly reduced consumption compared to unstressed controls (SMD -1.46; 95% CI: -1.65, -1.27; p < 0.0001). High variation in the results was still observed (between-study heterogeneity: I^2^ = 79.2%). Similarly, stressed rats treated with antidepressants had a significantly increased consumption compared to stressed controls (SMD 1.56; 95% CI: 1.23, 1.90; p < 0.0001). High between-study heterogeneity was observed (I^2^ = 74.6%).

##### *Bibliography*

1.	Zwetsloot, P. P. *et al.* Standardized mean differences cause funnel plot distortion in publication bias assessments. *Elife* **6**, e24260 (2017).

2.	Higgins, J. *et al.* *Cochrane Handbook for Systematic Reviews of Interventions | Cochrane Training*. (2022).

3.	Harrer, M., Cuijpers, P., Furukawa, T. & Ebert, D. dmetar: Companion R package for the guide ‘Doing Meta-Analysis in R’ — dmetar • dmetar. (2019).
